# Structure of an endogenous mycobacterial MCE lipid transporter

**DOI:** 10.1101/2022.12.08.519548

**Authors:** James Chen, Alice Fruhauf, Catherine Fan, Jackeline Ponce, Beatrix Ueberheide, Gira Bhabha, Damian C. Ekiert

**Affiliations:** Department of Cell Biology, New York University School of Medicine, New York, NY 10016, USA; Proteomics Laboratory, Division of Advanced Research Technologies, New York University School of Medicine, New York, NY 10016, USA; Department of Biochemistry and Molecular Pharmacology, NYU School of Medicine, New York, NY 10016, USA; Department of Neurology, New York University School of Medicine, New York, NY 10016, USA; Department of Microbiology, New York University School of Medicine, New York, NY 10016, USA

## Abstract

To replicate inside human macrophages and cause the disease tuberculosis, *Mycobacterium tuberculosis* (*Mtb*) must scavenge a variety of nutrients from the host^1,2^. The Mammalian Cell Entry (MCE) proteins are important virulence factors in *Mtb*^1,3^, where they are encoded in large gene clusters and have been implicated in the transport of fatty acids^4–7^ and cholesterol^1,4,8^ across the impermeable mycobacterial cell envelope. Very little is known about how cargos are transported across this barrier, and how the ~10 proteins encoded in a mycobacterial *mce* gene cluster might assemble to transport cargo across the cell envelope remains unknown. Here we report the cryo-EM structure of the endogenous Mce1 fatty acid import machine from *Mycobacterium smegmatis*, a non-pathogenic relative of *Mtb*. The structure reveals how the proteins of the Mce1 system assemble to form an elongated ABC transporter complex, long enough to span the cell envelope. The Mce1 complex is dominated by a curved, needle-like domain that appears to be unrelated to previously described protein structures, and creates a protected hydrophobic pathway for lipid transport across the periplasm. Unexpectedly, our structural data revealed the presence of a previously unknown subunit of the Mce1 complex, which we identified using a combination of cryo-EM and AlphaFold2, and name LucB. Our data lead to a structural model for Mce1-mediated fatty acid import across the mycobacterial cell envelope.

## Introduction

*Mycobacterium tuberculosis* (*Mtb*), the causative agent of tuberculosis, is one of the leading causes of death due to infectious disease, resulting in over one million deaths annually^9^. *Mtb* establishes a niche within the phagosomal compartment of host macrophages, where it can grow and replicate. To survive in the phagosome, *Mtb* must scavenge nutrients from the host cell^1,2^, and utilizes an ensemble of active transporters to import iron^10,11^, lipids^1,2^, and other metabolites^12^. In particular, the Mammalian Cell Entry (MCE) family of proteins has been implicated in the import of substrates such as fatty acids^4–7^ and cholesterol^1,4,8^ across the cell envelope of *Mtb* and related species such as *Mycobacterium smegmatis* (*Msmeg*) (Fig. 1a)^3,13,14^. MCE proteins are critical for virulence in *Mtb* and other bacterial pathogens^1,3,15–19^, underscoring their fundamental importance for nutrient acquisition from the host. To mediate the uptake of fatty acids and cholesterol, MCE transporters must translocate substrates across the impenetrable cell envelope, which consists of: 1) the inner membrane (IM), 2) the complex mycobacterial outer membrane (MOM), and 3) a periplasmic space between the IM and MOM, containing the cell wall^20^. In Gram-negative bacteria, many cargos are transported via large transenvelope proteinbased machines that mediate the passage of substrates across membranes and the periplasmic space, such as the LPS export system^21–26^, antibiotic efflux pumps^27,28^, and a variety of specialized protein secretion systems^29^. In contrast, it is unclear how substrates are transported across the highly divergent mycobacterial cell envelope, whether such periplasm-spanning complexes exist, and how active transporters such as the MCE transporters may facilitate substrate transport in mycobacteria.

**Fig. 1:**
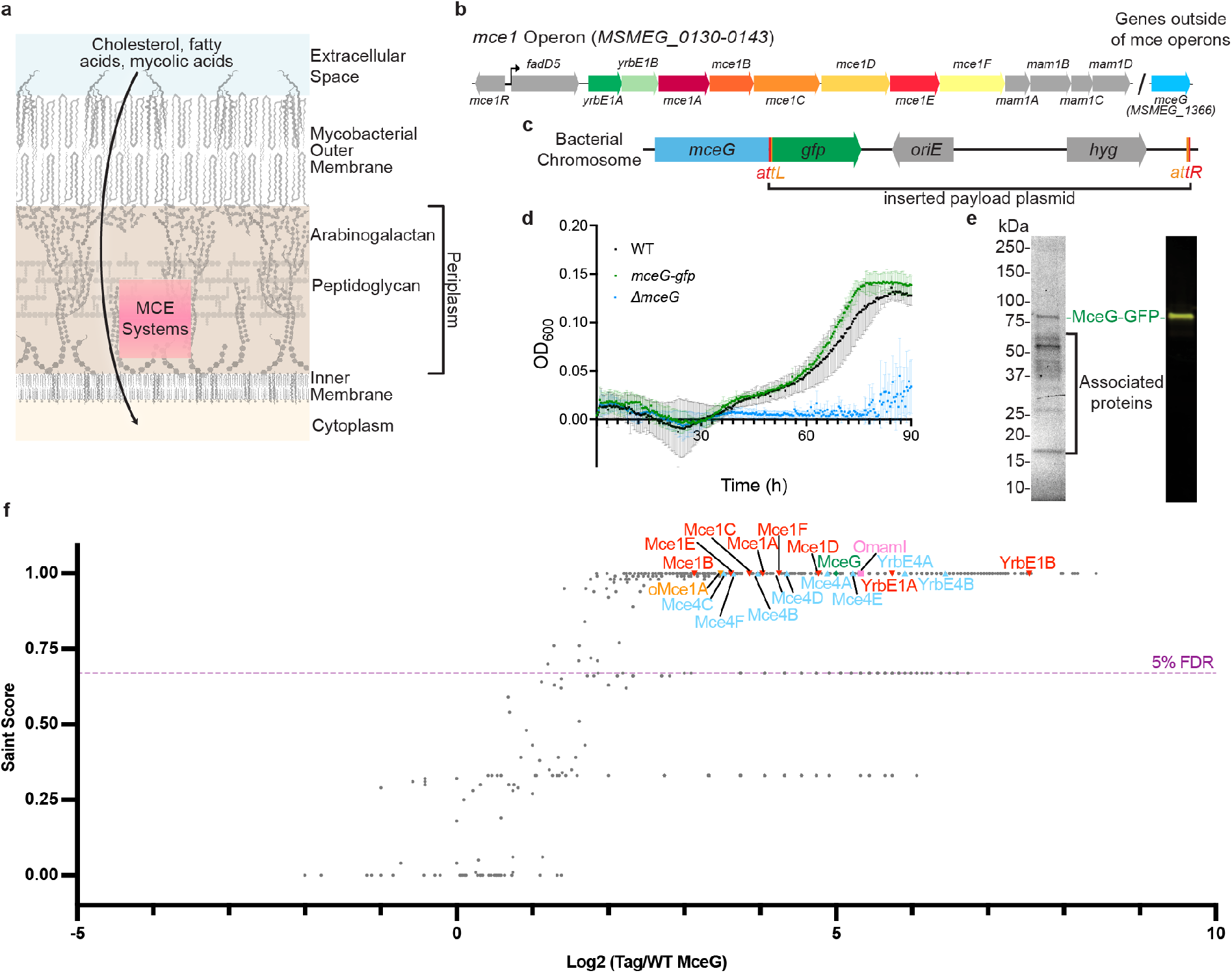
Construction and purification of MceG-GFP. **a**, Schematic of mycobacterial cell envelope, adapted from Dulberger et al.^20^. MCE systems are proposed to facilitate transport of nutrients across the cell envelope. **b**, Schematic of *Mycobacterium smegmatis (Msmeg) mce1* operon. **c**, Schematic of *Msmeg* bacterial chromosome modified by ORBIT^37^. The gene *mceG (MSMEG_1366)* is shown with inserted payload plasmid containing *gfp, oriE*, and *hyg* flanked by an *attL* site and an *attR* site. **d**, Growth curve of *mceG-gfp* strain (green) compared to Wild-Type *Msmeg* mc^2^155 strain (WT, black) and *ΔmceG* (blue) in minimal media containing cholesterol. Y-axis shows the optical density at 600 nm for bacterial cultures and X-axis shows incubation time in hours. Growth assays were repeated three times (*n* = 3) with similar results. Plotted data are the mean of three replicates and standard error bars are shown. **e**, (left) Stain-free SDS-PAGE of MceG-GFP pulldown in n-dodecyl-β-D-maltoside (DDM) after GFP-affinity purification and size exclusion chromatography. (right) Corresponding Western blot using an anti-GFP antibody against purified MceG-GFP. **f**, Plot of proteins identified by mass spectrometry that co-purify with MceG-GFP. Each point corresponds to an individual protein plotted by fold change difference after purification of MceG-GFP from *Msmeg* strain harboring tagged MceG versus control wild-type *Msmeg* mc^2^155 (x-axis) and the probability that a protein is a MceG interactor (SAINT score; y-axis). SAINT score = 1 identifies proteins with the highest probability of being a MceG interactor^59^. SAINT score ≥ 0.67 yielded an FDR (false discovery rate) of ≤ 5% as indicated by the purple dotted line. Proteins related to the mycobacterial MCE systems are highlighted and annotated: Mce1 proteins (red upside-down triangles), Mce4 proteins (sky blue triangles), orphaned MCE proteins (orange upside-down triangle), orphaned Mce-associated membrane proteins (pink square), MceG (green diamond). Proteins outside of these systems are shown as grey dots. Plotted data are from three biological replicates (*n* = 3).

In *Mtb*, MCE transport systems are encoded in four different gene clusters, *mce1-mce4*, which are among the largest operons in the genome (Extended Data Fig. 1a). Each cluster has a core module of eight conserved genes: 1) two *yrbE* genes encoding the transmembrane subunits of an ATP-binding-cassette (ABC) transporter and 2) six genes encoding MCE proteins. A variable number of “accessory” proteins are often found adjacent to the eight-gene core module^13^. Additional proteins encoded elsewhere in the genome are also required for *Mtb* MCE transporter function, including an ATPase, MceG^1,30^, and an integral membrane protein, LucA^4,5^. This gene organization is conserved in other mycobacterial species, including *Msmeg* (Fig. 1b, Extended Data Fig. 1b)^31,32^, and the proteins from each gene cluster are thought to interact with each other to form large complexes^14^. Recombinant expression and purification of MCE complexes has been challenging due to the complexity of their genetic organization, and studies thus far have been limited to single subunits and smaller subcomplexes^33,34^. Thus, how proteins are arranged in a complex to facilitate lipid transport across the cell envelope remains unclear, and elucidating the architecture of mycobacterial MCE systems is a key step towards understanding their transport mechanism.

## Results

### Isolating endogenous MCE complexes from Mycobacterium smegmatis

To isolate intact complexes for structural studies in the absence of an established recombinant expression system, we purified endogenous MCE transporters from *Msmeg*, which have high sequence identity to their *Mtb* orthologs (~68 % identical^31^) and similar functions^8,34–36^. We inserted a GFP tag at the C-terminus of MceG in the chromosome of *M. smegmatis mc^2^155* using homologous recombination via ORBIT (Fig. 1c and Supplementary Table 1)^37^. Tagging the C-terminus of MceG did not significantly impact growth using cholesterol as the sole carbon source in an established assay^14^, indicating that the MceG-GFP fusion is functional (Fig. 1d). The GFP tag on MceG was used for affinity purification of endogenous MCE complexes from *Msmeg* cells (Fig. 1e and Extended Data Fig. 1c). Because MceG is thought to be shared between multiple MCE systems in a given bacterial species^30,35^, pulling down MceG-GFP may lead to the purification of a mixture of several MCE complexes expressed in *Msmeg* under our experimental conditions. To identify the protein subunits that form complexes with MceG and to assess the complexity of our sample, we used mass spectrometry. These experiments revealed that MceG co-purifies with the eight core components from each of the *mce1* and *mce4* operons, including both YrbEs and all 6 MCE proteins (Fig. 1f, Supplementary Tables 2,3). Mce1 has been shown to transport fatty acids and mycolic acids^4–7^, whereas Mce4 imports cholesterol^1,4,8^. Quantification of relative protein abundance based on peptide spectral matches shows that Mce1 subunits are most abundant (Supplementary Table 2). We did not observe any peptides corresponding to *mce1*-encoded proteins Mce1R, FadD5, or Mam1A-Mam1D, or the accessory factor LucA. MSMEG_6540, which is 84% identical to Mce1A, but encoded elsewhere in the genome, was also highly enriched in the MceG pull-down and has recently been proposed to play a role in Mce1-mediated fatty acid uptake^34^. While most other mycobacterial MCE proteins are encoded in 6-gene modules, MSMEG_6540 is an “<u>o</u>rphaned” paralog of Mce1A found in a single gene operon, which we therefore name oMce1A.

### Overall structure of the Mce1 transporter

We determined the structure of the Mce1 transporter using single-particle cryo-electron microscopy (cryo-EM) (Extended Data Figs. 2a-c, 3a) to a resolution ranging from ~2.30 Å to ~3.20 Å (Map0, Fig. 2a, Extended Data Figs. 3b-d, Supplementary Table 4). While our mass spectrometry data indicate a mixture of Mce1 and Mce4 in the sample used for cryo-EM, side chain density throughout our final high-resolution map unambiguously shows that our map corresponds to the Mce1 complex (Extended Data Fig. 4a), and we do not see any evidence of Mce4 subunits (see Methods). The Mce1 complex consists of 10 protein subunits, including two copies of MceG and a single copy each of YrbE1A, YrbE1B, Mce1A/oMce1A, Mce1B, Mce1C, Mce1D, Mce1E, and Mce1F (Fig. 2b). Several proteins encoded in the *mce1* operon were absent from the complex, including FadD5 and Mam1A-Mam1D, suggesting that they may bind with lower affinity, transiently, or may not interact directly. Density for the Mce1A subunit is ambiguous at residues that differ between Mce1A and oMce1A, suggesting that our reconstruction contains a mixture of these highly homologous proteins at the location of the Mce1A subunit (see Methods). Our final model is nearly complete, apart from regions predicted to be unstructured near the C-termini of Mce1C, Mce1D, and Mce1F (Extended Data Fig. 4d).

**Fig. 2:**
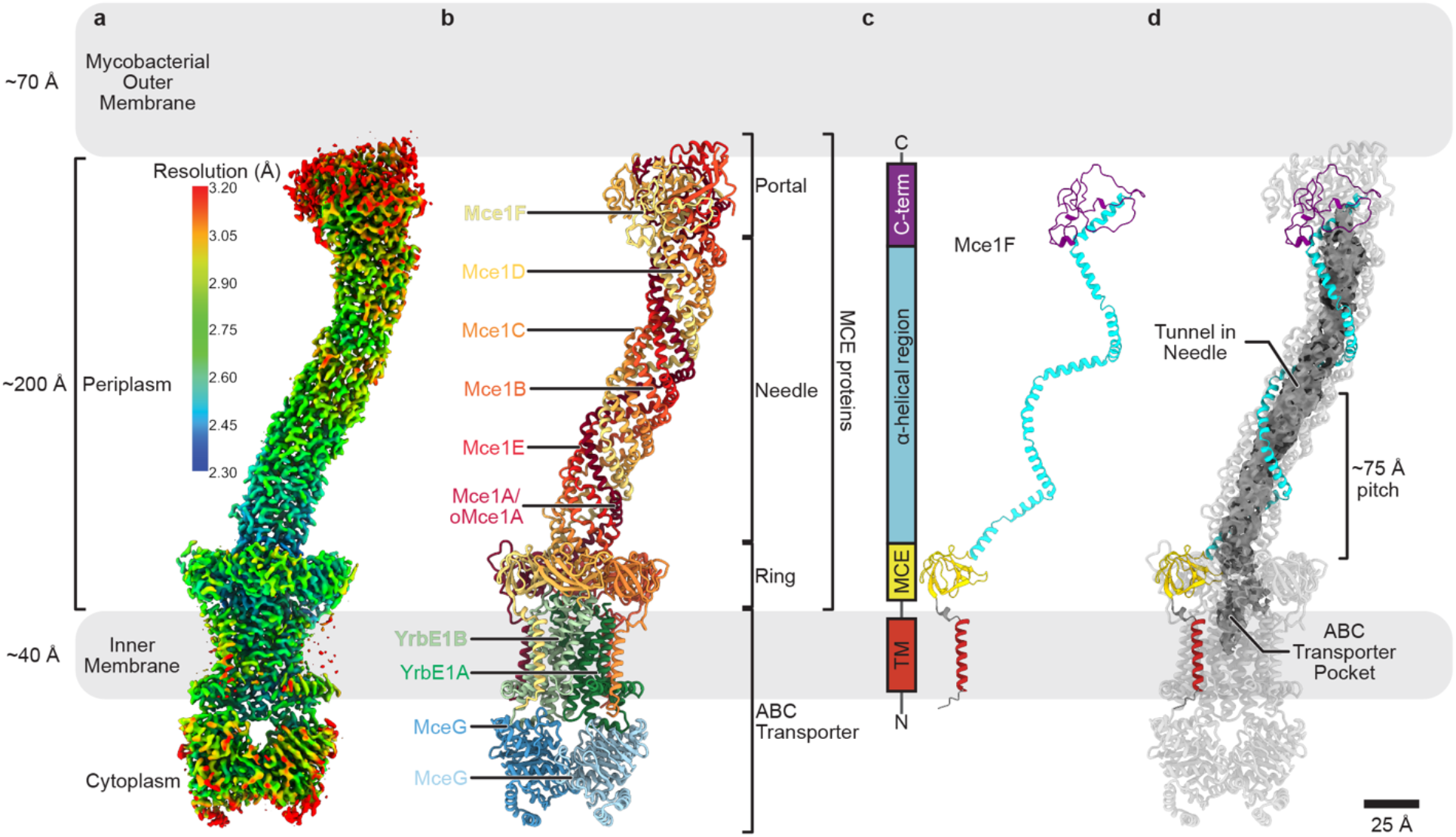
Cryo-EM structure of an endogenous MCE complex. **a-d**, Simplified diagram of the mycobacterial cell envelope drawn to scale. **a**, Composite cryo-EM map (Map0) of complex from MceG-GFP pulldown colored by local resolution as estimated using cryoSPARC^60^. **b**, Structure of Mce1 transport system, corresponding to the map shown in Fig. 2a and colored by subunit as in Fig. 1b. The four main parts of the structure are labeled: portal, needle, ring and ABC transporter. The portal, needle and ring are formed from different regions of the six MCE proteins. **c**, Mce1F extracted from Fig. 2b to highlight the structure of an individual MCE protomer. Colors are consistent in the 2D schematic and 3D structure, showing different regions of the MCE protein. **d**, Mce1 structure shown as a cartoon with the tunnel within the needle assembly and substrate-binding pocket in the ABC transporter rendered as a molecular surface and colored grey (calculated in CASTp 3.0^61^). Mce1F is colored as in Fig. 2c.

Mce1 forms a highly elongated complex, ~310 Å in length, which can be divided into four main parts (Figs. 2b,c, Supplementary Video 1): 1) the portal, a globular domain formed by the C-termini of the Mce1ABCDEF subunits, that lies proximal to the MOM; 2) the needle, which consists of a long central tunnel and is formed by the α**-** helical regions of the Mce1ABCDEF subunits; 3) the ring, formed by the MCE domains of the Mce1ABCDEF subunits; and 4) the ABC transporter in the IM, which consists of YrbE1AB permease subunits and MceG ATPase subunits. The Mce1 complex is anchored in the IM at one end, and the portal, needle, and ring extend ~225 Å into the periplasmic space. As the periplasmic width of *Msmeg* is ~200 Å^38^, the Mce1 complex is long enough to span the distance between the MOM and IM, with the potential to import fatty acids through its central tunnel, shielded from the surrounding hydrophilic space (Fig. 2d). This is conceptually similar to molecular machines in Gram-negative bacteria that form tunnels and bridges to move small hydrophobic molecules across the periplasm. However, the elongated tunnel of Mce1 is structurally divergent from proteins characterized to date (Extended Data Fig. 5a), and to our knowledge in the first structure of such a periplasm-spanning transport system in mycobacteria (Extended Data Fig. 5b).

### The portal creates an entrance to the transport pathway

Substrates for import from the MOM may enter the Mce1 complex through the portal domain (Fig. 3a), which is composed of a small six-stranded β-barrel (Fig. 3b) surrounded by non-canonically structured regions (Extended Data Fig. 6a,b). Apart from the β-barrel motif, the portal domain has no apparent homology to any known protein domains. The C-terminus of each MCE protein contributes a single β-strand to the formation of the β-barrel, and also provides a portion of the surrounding non-canonical regions. Despite being formed from six homologous MCE proteins (Mce1A-Mce1F), the C-terminal regions of each MCE subunit are structurally distinct and vary widely in length (Extended Data Fig. 6a,b). The lumen of the β-barrel is aligned with the tunnel and has a hydrophobic interior, potentially acting as an entry point for substrates (Fig. 3c). While this β-barrel is formed from just 6 strands, the high tilt of its β-strands results in a barrel diameter similar to the 8-stranded fatty acid binding phospholipase PagP found in the *E. coli* outer membrane^39^. In our structure, passage through the β-barrel is blocked by a few loosely packed hydrophobic side chains that protrude into the lumen. If and how opening may occur is unclear, but relatively subtle side chain rearrangements may be sufficient to open a pore large enough for a fatty acid to thread through.

**Fig. 3.**
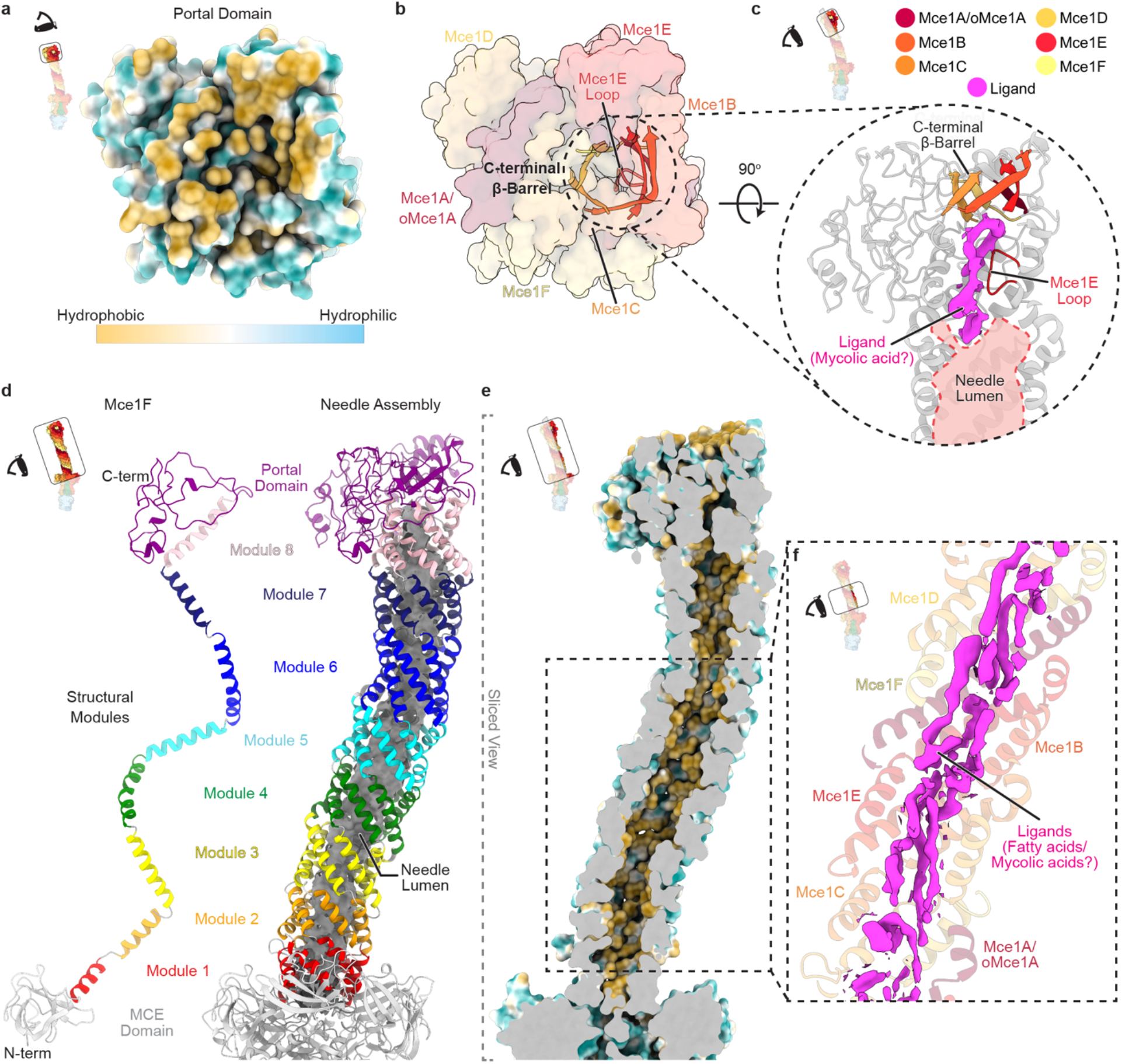
Architecture of portal and needle domains of the Mce1 system. **a**, View of the portal domain from the outer membrane. The inset highlights the region being shown and the schematic of the eye designates the point of view. Proteins are shown as a molecular surface and colored by hydrophobicity in ChimeraX^62^. **b**, Same view as Fig. 3a. Proteins are shown as a transparent molecular surface, and the C-terminal β-barrel and Mce1E loop are shown in cartoon representation with bright colors. **c**, Rotated zoomed-in view of the region circled in Fig. 3b as indicated by the inset on the top-left. The Mce1E loop and C-terminal β-barrel, are shown in color, and the surrounding protein regions are grey. Ligand density from Map0 colored magenta. The lumen of the needle is outlined and colored in light red. **d**, (left) One MCE protomer, Mce1F extracted from the hexamer. MCE needle modules are colored in rainbow colors (module 1, red to portal, purple) and MCE domains are light grey. (right) Mce1 needle domain, colored by module. The lumen of the needle is rendered as a grey molecular surface (calculated using CASTp 3.0^61^). **e**, Needle lining shown by a sliced view of Fig. 3d as indicated in the inset. Protein is rendered as a molecular surface and colored by hydrophobicity in ChimeraX ^62^. Color key is shown in Fig. 3a. **f**, Ligand density (magenta) from Map0 in the region of the needle boxed in Fig. 3e and indicated in the top-left inset.

### The needle forms a unique tunnel assembly to facilitate transport of substrates

The portal feeds directly into a tunnel created by the needle, a unique α-helical structure that is strikingly curved. Our EM data for Mce1 suggest that the curved needle is fairly rigid, and we do not observe straight or alternatively-curved states. The needle curvature likely arises from the asymmetric, heterohexameric assembly of the MCE proteins, but its functional role is not immediately clear. Each MCE subunit contains eight copies of a helical repeat motif, separated by well-defined kinks (Fig. 3d, Extended Data Fig. 6a). The helical segments from Mce1ABCDEF twist around each other to form a left-handed superhelix with a pitch of ~75 Å and almost exactly two complete turns (Fig. 2d). The first helical repeats from each MCE subunit associate to form a 6-helix bundle. Similarly, repeats 2, 3, 4, 5, 6, 7, and 8 associate to form separate 6-helix bundles, for a total of eight structurally similar modules (Extended Data Fig. 6c). These eight modules stack on top of each other to make a long, needle-like tube, and are connected by short linkers (Fig. 3d). The 6-helix bundles appear to be unrelated to previously described folds, such as 6-helix coiled-coils^40^.

The inside of the needle contains a long tunnel, ~7,000 Å^3^ in volume, with an inner diameter ranging from 7-11 Å. The tunnel is lined with hydrophobic residues, potentially providing a sheltered passageway for fatty acids to cross the periplasm (Fig. 3e, Extended Data Fig. 6d). Numerous strong densities are present in the needle, which may correspond to bound substrates (Fig. 3f). The resolution of these densities is too low to unambiguously identify the ligand, but the size and shape are consistent with fatty acid chains that range from 10 to 49 carbons in length (Extended Data Fig. 4b). In many places, 3-5 fatty acid-like densities appear to run parallel to each other along the long axis of the needle, suggesting that multiple substrates may be transported “in bulk” through the tunnel. One of the largest and most prominent densities is located in the needle just below the portal domain, where a loop from Mce1E protrudes into the lumen and partially occludes the otherwise broad and featureless tunnel (Figs. 3b,c). The constriction in the tunnel formed by this loop may create a fatty acid binding site reminiscent of the high affinity site in the long-chain fatty acid transporter, FadL^41^. In our structure, strong density for a possible mycolic acid substrate (49-carbons) fills the area surrounding this loop (Fig. 3c), consistent with a possible role of Mce1 in mycolic acid recycling and MOM maintenance^7^. This binding site, just beyond the β-barrel entrance, may be involved in substrate selection, occurring prior to transport through the tunnel.

### MCE ring connects needle to an ABC transporter

The hydrophobic tunnel through the needle leads to a pore through the ring, which is formed by six MCE domains (Fig. 4a). Each MCE domain in the ring is structurally similar (Extended Data Fig. 7a) but the domains are only ~17% identical to one another at the sequence level (Extended Data Fig. 7b), leading to a heterohexameric ring with the following arrangement: Mce1A/oMce1A-Mce1E-Mce1B-Mce1C-Mce1D-Mce1F (Fig. 4b). This contrasts with the rings observed in other MCE protein assemblies, including LetB, PqiB, and MlaD, which are homohexameric and approximately six-fold symmetric^42,43^. The pore of the Mce1 ring is formed by a pore-lining loop (PLL) from each MCE domain (Fig. 4b, Extended Data Fig. 7c). The arrangement of the PLLs may form a gate between the periplasmic needle assembly and the substrate-binding pocket of the ABC transporter below (Fig. 4c). In our structure, the pore through the ring is closed, and a conformational change is likely required to allow passage of substrates into the ABC transporter. Opening and closing of the tunnel through MCE rings has been observed previously in LetB and PqiB^42,43^, and may also occur in the Mce1 ring.

**Fig. 4:**
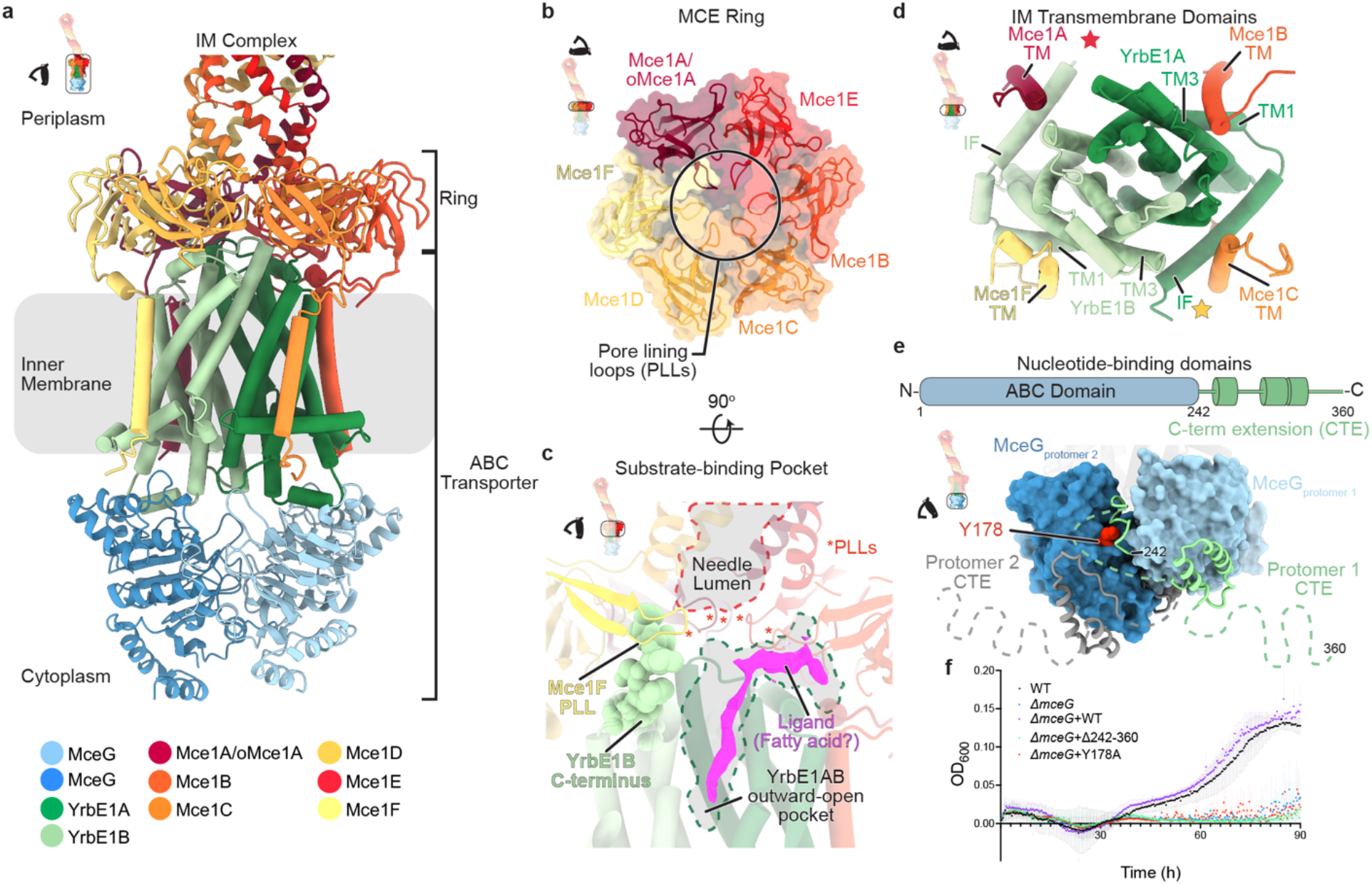
Architecture of the ring and ABC transporter complex at the inner membrane. **a**, Inner membrane complex of Mce1, including the ring and ABC transporter as indicated in inset. Structure is colored according to the key below. **b**, Mce1ABCDEF heterohexameric MCE ring viewed from the periplasm, as indicated by inset. Proteins colored according to the key in Fig. 4a. MCE domains of Mce1ABCDEF are shown as cartoon representations superimposed on a transparent molecular surface. **c**, Interface between the Mce1ABCDEF heterohexameric MCE ring and YrbE1AB heterodimer. Proteins colored according to the key in Fig. 4a and pore-lining loops (PLLs) are indicated with red asterisks. The C-terminus of YrbE1B is shown with spheres and interacts with Mce1F PLL. The needle lumen and ABC transporter pocket are outlined with dotted lines. Ligand density in the YrbE1AB cavity from Map0 is colored magenta. **d**, ABC transporter transmembrane domains composed of YrbE1AB and the interacting Mce1ABCDEF transmembrane helices (TMs), as indicated in top-left inset. Helices are shown as cylinders and colored according to Fig. 4a. Colored stars represent the expected position of Mce1E lipid anchor and Mce1D TM based on structural alignment with *E. coli* MlaFEDB (PDB ID 6XBD)^46^. These two regions were not resolved in our map. **e**, (top) Domain schematic of MceG designating residue boundaries for ABC domain and C-terminal extension. (bottom) MceG ATPase subunit homodimer as indicated by inset. ABC domains are shown as blue molecular surfaces, and the C-terminal extension is shown as green (MceG_protomer 1_) or grey (MceG_protomer 2_) cartoons. Regions that were not modeled in the cryo-EM map due to unresolved density are indicated by dotted lines. MceGprotomer 2 residue (Y178) that interacts with the C-terminal extension of MceG_protomer 1_ is shown as red spheres. **f**, Cholesterol growth curves of *ΔmceG* strain complemented with plasmids containing the following *Msmeg* MceG mutants: 1) MceG Δ294-360 (pale green) and 2) MceG Y178A (red). Wild-type *Msmeg* mc^2^155 strain (WT, black), Δ*mceG* (blue), and Δ*mceG* complemented with wild-type *mceG* (Δ*mceG*+comp, purple) are shown as controls. Growth assays were repeated three times (*n* = 3) with similar results. Plotted data are the mean of three replicates and standard error bars are shown.

### ABC transporter in the inner membrane is poised to accept substrates from MCE ring

The pore through the MCE ring leads to the ABC transporter in the IM, which consists of a heterodimer of permease proteins, YrbE1A and YrbE1B and a homodimer of the ATPase MceG (Fig. 4a). YrbE1A and YrbE1B each consist of an N-terminal interfacial helix and five TM helices, and are homologous to the transmembrane domains of the recently described type VIII ABC transporter, MlaFEDB (Extended Data Figs. 8a,b) ^44–46^. The TMs of Mce1A, B, C, and F are well resolved and clearly interact around the periphery of the ABC transporter transmembrane domains and anchor the MCE ring in place (Fig. 4d). The TM helix of Mce1D and lipid anchor of lipoprotein Mce1E are not resolved in our structure but may also play similar roles. The MCE ring is slightly tilted with respect to the YrbE subunits (~4°) (Extended Data Fig. 8c), reminiscent of conformations previously described in the homologous MlaFEDB MCE transporter from *E. coli*^46^. The C-terminus of YrbE1B wedges into the space between the MCE ring and the YrbEs, making contacts with the Mce1F PLL (Fig. 4c, Extended Data Fig. 8d). This extension may stabilize the tilted state, possibly playing a role in coupling conformational changes in the ABC transporter to MCE ring opening/closing. In contrast to the homodimer found in most bacterial ABC transporters, the YrbE1AB heterodimer could facilitate the recognition of asymmetric substrates^47^.

In our structure, YrbE1AB adopts an outward-open state, with a narrow substrate-binding pocket of ~150 Å^3^ that is formed between the YrbE subunits (Figs. 2d,4c). Density for an elongated ligand, resembling a fatty acid, is observed extending upwards from the substrate binding pocket (Fig. 4c). An MceG ATPase is bound to each YrbE subunit, forming a homodimer (Fig. 4e). Each MceG contains a ~120 amino acid C-terminal extension that is much longer than canonical ABC transporters. This extension consists of several α-helices connected by flexible linkers that interact with the neighboring MceG subunit (Fig. 4e). Cholesterol growth assays with MceG mutants demonstrate that the C-terminal extension and its interaction with the neighboring subunit is important for function (Fig. 4f), consistent with previous findings^35^. Our results suggest that the extension may be important for stabilizing the MceG homodimer, as recently proposed for another MCE transporter^48^, or may play a regulatory role in ATP hydrolysis or substrate transport. No significant density was observed in the MceG ATP-binding site and the dimer is open, allowing nucleotide exchange. Our structure suggests that the resting state of the Mce1 complex is outward-open, similar to the MlaFEDB phospholipid transporter^46,49–53^ and the LptBFG LPS transporter^54,55^.

### LucB is a novel subunit of the Mce1 transporter

Unexpectedly, we observed density for an additional unknown subunit associated with the ABC transporter within a subpopulation of our particles (Extended Data Fig. 2a). Focused 3D classification led to the emergence of two classes (Fig. 5a), Class 1 (Map1, ~2.76 Å, Extended Data Figs. 3e-h, Supplementary Table 4) and Class 2 (Map2, ~2.90 Å, Extended Data Figs. 3i-l, Supplementary Table 4). The additional subunit, found only in Class 1, lies almost entirely within the transmembrane region, and consists of 4 TM helices (Fig. 5b). Examination of our MceG-GFP mass spectrometry data did not suggest an obvious candidate protein consistent with our EM density (Supplementary Tables 2,3). To identify this unknown subunit, we built a polyalanine model into the density and used these coordinates to do a structure-based search of the Protein Data Bank and AlphaFold Protein Structure Database^56^ using Foldseek (Fig. 5b)^57^. While no proteins with similar structure were identified in the Protein Data Bank, the search of the AlphaFold database revealed predicted structures that matched our polyalanine model well, including MSMEG_3032 and its *Mtb* homolog Rv2536^58^ (~61% identical) (Fig. 5c). Fitting the AlphaFold2 MSMEG_3032 model into our EM density required minimal adjustment apart from a few sidechain rotamer changes, supporting the assignment of MSMEG_3032/Rv2536 as a novel component of the Mce1 system (Fig. 5d, Extended Data Fig. 4c). Based upon a possible role as a <u>L</u>ipid <u>U</u>ptake <u>C</u>oordinator, analogous to the proposed role of LucA^4^, we rename MSMEG_3032/Rv2536 to LucB. To validate the interaction identified from our structure, we assessed whether LucB pulled down MCE transporter components. We constructed an *Msmeg* strain with chromosomally tagged LucB-GFP, and purified the protein by anti-GFP affinity and size exclusion chromatography (Extended Data Figs. 9a,b).

**Fig. 5:**
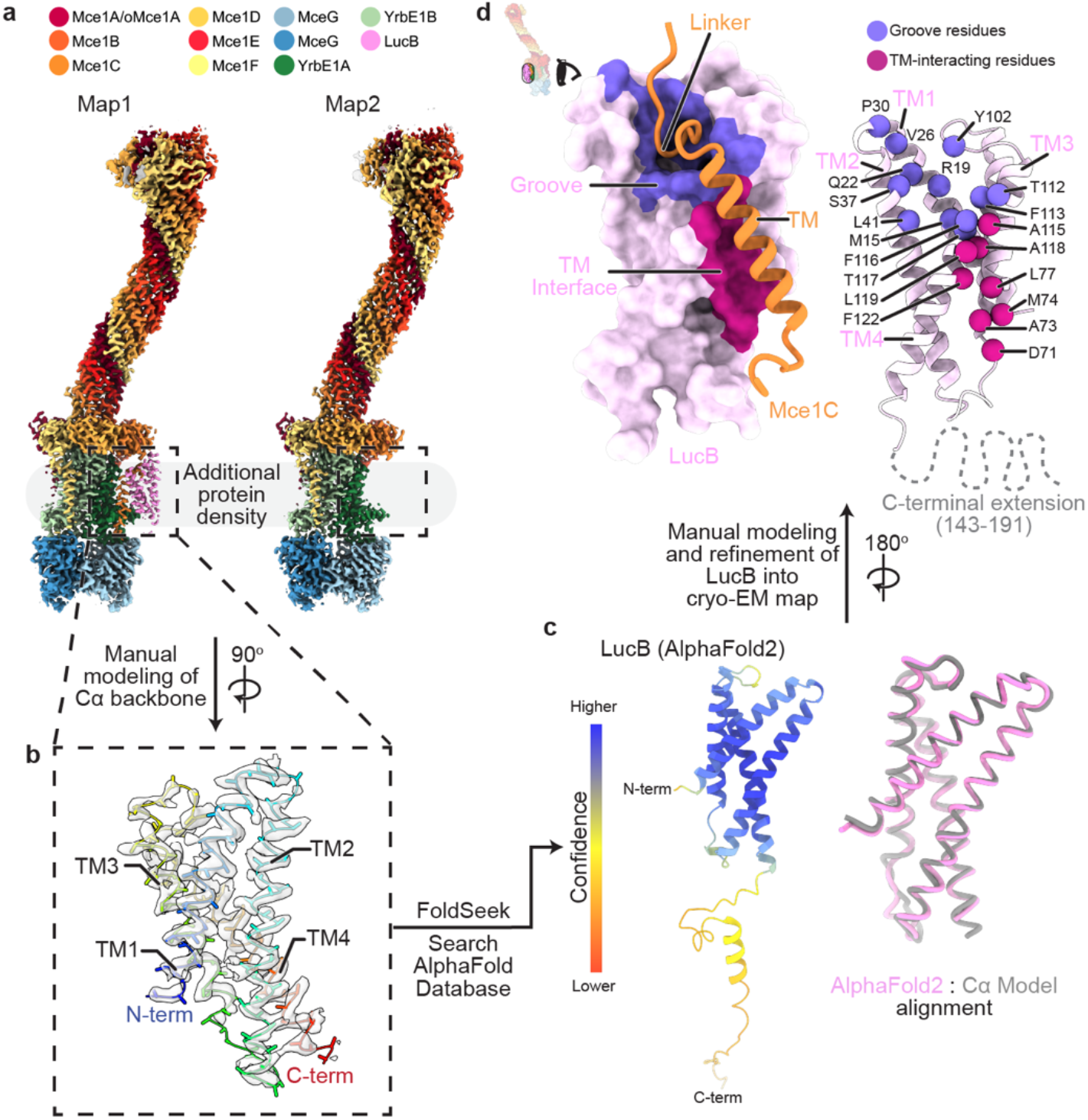
LucB is an accessory factor that binds the Mce1 complex. **a-d**, Workflow to identify unknown protein subunit for which density is observed in Class 1 (Map1). **a**, Composite density maps for Class 1 (Map1) and Class 2 (Map2) colored by protein subunit. Color key is shown above. Pink density is observed only in Class 1. **b**, Cα backbone manually built into extra density observed in Class 1. 3D model of poly-alanine Cα backbone was used as a search model for Foldseek^57^. Protein is colored in rainbow colors (N-terminus, blue; C-terminus, red), and corresponding cryo-EM density is shown as a transparent grey surface. **c**, (left) AlphaFold2 prediction of LucB identified from Foldseek^57^ search. Model is colored by prediction confidence; the N-terminal domain is predicted with high confidence^63^. (right) Structural alignment of Cα backbone and LucB AlphaFold2 prediction for the N-terminal domain. **d**, Mode of interaction between LucB and the Mce1 complex as indicated by inset. (left) LucB (pink) is rendered as a molecular surface and Mce1C (orange) is shown in cartoon representation. LucB binding interfaces are colored: groove (purple) and TM interface (violet). (right) LucB and Mce1C are shown as cartoons. Residues that make up the LucB groove and TM interface are depicted as spheres and color as left.

Negative stain electron microscopy of the resulting sample reveals particles with characteristic shape and features of the Mce1 system (Extended Data Figs. 9c,d). Mass spectrometry of purified LucB-GFP (Extended Data Fig. 9e, Supplementary Tables 2,5) showed significant enrichment of Mce1 subunits, while Mce4 subunits were not significantly enriched. Together, these data suggest that LucB preferentially associates with the Mce1 transporter under our experimental conditions.

In our structure, LucB interacts almost exclusively with Mce1C, primarily via interactions with the Mce1C TM helix and linker connecting the TM helix to the MCE domain (Fig. 5d). The Mce1C linker sits in a conserved cleft formed between the TM2 and TM4 helices of LucB (Extended Data Fig. 10a), and the Mce1C TM packs against TM3 and TM4 of LucB. Binding to Mce1C positions the LucB C-terminal extension towards the cytoplasm where it could potentially interact with MceG or recruit other proteins (Fig. 5d). The C-terminal extension is not resolved in our map and is predicted to be disordered (Fig. 5c), but may become ordered upon interacting with a binding partner. The conformation of the Mce1 complex is very similar in both classes, apart from clear definition of density for the Mce1C transmembrane helix and interacting loop in the presence of LucB (overall RMSD = 0.50 for Class 1 Vs. Class 2), suggesting that there is no global conformational change in the Mce1 system upon LucB binding.

LucB, for which there is a single paralog in *Msmeg* and *Mtb*, is a protein of unknown function and has not previously been linked to MCE transporters. Orthologs of this protein can be found in bacteria of the Actinomycetales order, particularly in the families: Gordoniaceae, Mycobacteriaceae, Nocardiaceae, Pseudonocardiaceae, and Tsukamurellaceae (Extended Data Fig. 10b). Interestingly, LucB orthologs appear to be found only in double-membraned bacteria containing *Mtb*-like *mce* operons^8^, with a conserved 8 gene cluster encoding two distinct YrbE and six distinct MCE proteins. Conversely, orthologs of LucB are not found in genomes that encode simpler MCE gene clusters encoding single YrbE and MCE proteins subunits, such as those found in *E. coli*. This observation, coupled with our data, suggests that LucB may have evolved to function specifically with heterooligomeric MCE transporters that arose in the actinobacterial lineage, and may be involved in the regulation of activity in these transporters.

## Discussion

The mycobacterial cell envelope is highly complex and divergent from its Gram-negative counterparts. Mechanisms for how substrates are transported across the mycobacterial cell envelope have remained elusive. Our high-resolution structure of an endogenous Mce1 transport complex allows us to propose a model for how this important virulence factor may work to import substrates (Fig. 6, Supplementary Video 2). First, fatty acids or mycolic acids from the MOM may enter through the β-barrel of the portal domain, either directly or mediated by additional unknown factors in the MOM. How the Mce1 complex recognizes specific substrates is unclear, but one possibility is that substrate selection occurs at the apparent fatty acid binding site noted just below the β-barrel of the portal domain. After entering the complex, the substrates travel across the periplasm through the hydrophobic tunnel created by the curved Mce1ABCDEF needle, in which several substrates may be accommodated simultaneously. At the base of this needle, the ring of MCE domains must undergo a conformational change, opening the central pore to allow substrate entry into the IM ABC transporter. ATP hydrolysis by MceG likely drives conformational changes in the YrbE1AB subunits to translocate substrates into the cytoplasm or IM. LucB, which we show binds to Mce1C, may play a role as a regulator, or a scaffold protein to recruit other parts of the system that are not yet known. While LucB is not structurally related to LucA, both are small transmembrane proteins that may regulate MCE systems. Our data provide a structural framework for how mycobacteria may use MCE systems to scavenge resources, such as fatty acids, from the host cell by providing a tunnel for the transport of substrates across the cell envelope without compromising the protective nature of this barrier.

**Fig. 6:**
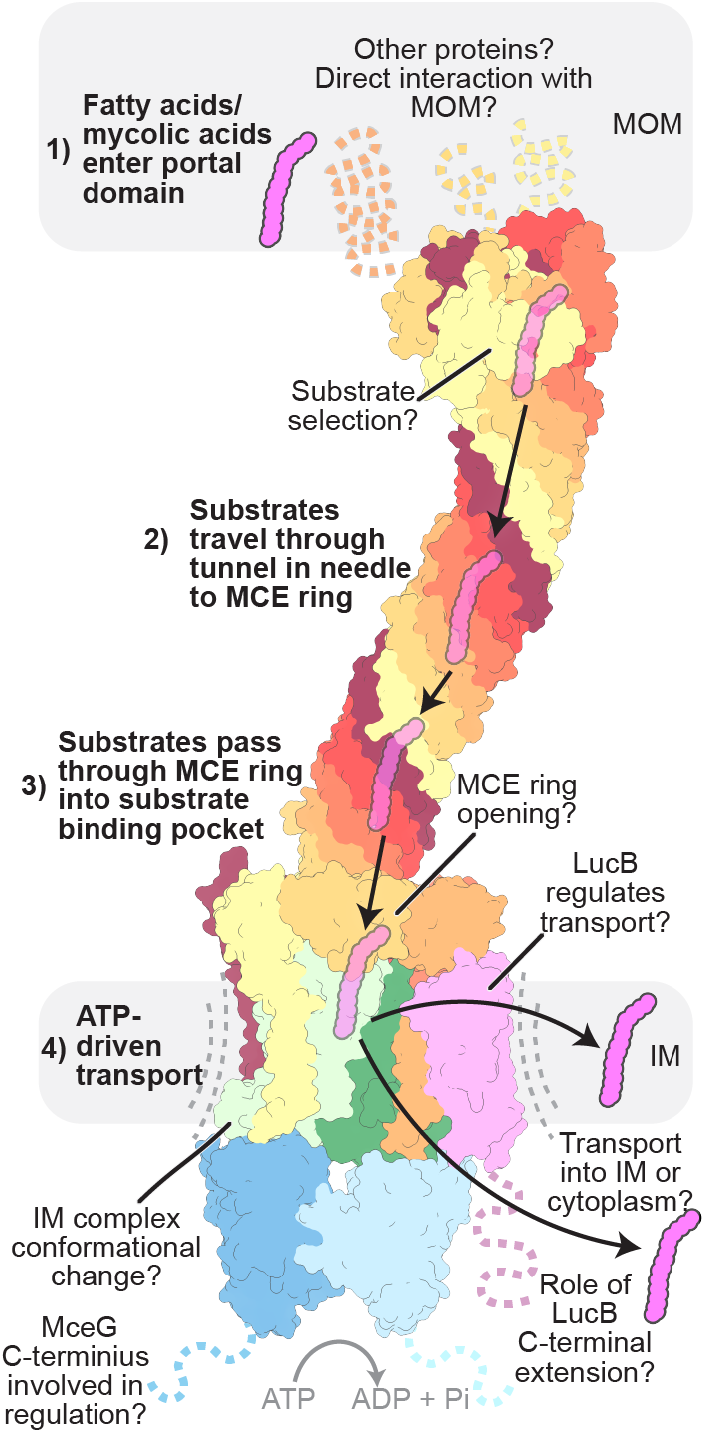
Model for Mce1-mediated transport. Model for Mce1-mediated transport of fatty acids in mycobacteria, highlighting our current understanding and open questions: 1) fatty acids (magenta) enter the portal domain; 2) substrates travel down the hydrophobic tunnel through the needle domain; 3) substrates pass through the ring into the ABC transporter; 4) ATP hydrolysis in MceG drives conformational changes in the ABC transporter, allowing substrates to be transported. The nature of the conformational changes that drive transport and how LucB regulates Mce1 remain unknown.

## Methods

No statistical methods were used to predetermine sample size. The experiments were not randomized, and the investigators were not blinded to allocation during experiments and outcome assessment.

### Bacterial strain construction

*Mycobacterium smegmatis* (*Msmeg*) strains were generated by the oligonucleotide-mediated recombineering followed by Bxb integrase targeting (ORBIT)^37^. An expression plasmid (pKM444, Addgene #108319, for tagging or pKM461, Addgene #108320, for knockouts)^37^ containing the Che9c phage RecT annealase and Bxb1 integrase was electroporated into electrocompetent *Msmeg* cells (mc^2^155 strain^73^) and protein expression was induced with 500 ng/mL anhydrotetracycline (ATc, Sigma, cat. #31741). For chromosomal tagging, the induced cells were made electrocompetent and subsequently co-transformed with pBEL2108 (a derivative of payload plasmid pKM468 (Addgene #108434)^37^ containing a 3C protease cleavage site upstream of the eGFP tag) and a targeting oligonucleotide. MceG-GFP strain (bBEL591) was generated with a 3C-eGFP-4xGly-TEV-Flag-6xHis tag on the C-terminus of MceG (*MSMEG_1366*) using the following oligo (IDT Ultramer DNA Oligo): 5’-GTTGCCCGCGCGCCGGCCCCTTGAGACACGTCAGGCCGGGCCGTGACGGCCCGGCCTGATCGCGGCAAACTCAGGTTTGTACCGTACACCACTGAGACCGCGGTGGTTGACCAGACAAACCCGCCTGCTTGGGCACCTCGATGACGCCCGTCGGCGAGTCGTCGTAGTTCTCGACGGGCGCGGTGGCGGCC-3’. LucB-GFP (bBEL595) strain was generated with a 3C-eGFP-4xGly-TEV-Flag-6xHis tag on the C-terminus of LucB (*MSMEG_3032)* using the following oligo (IDT Ultramer DNA Oligo): 5’-CACGATGTGTGACGCTACTCGCTACGCTGTGCCCCCATGAGCAAGTGGTTACTGCGCGGAGTGGTGTTCGCAGGTTTGTCTGGTCAACCACCGCGGTCTCAGTGGTGTACGGTACAAACCCCGCTGGAGAATCCGGACCAGCCGCGTCAGAGCTGATCCGGGCTCAGCTTCACAAACGAGAGTTGTTGTGGT-3’.

Transformants were plated on either LB+agar (Luria-Bertanior, Difco cat. #DF0446-07-5) or 7H10 (Difco, cat# DF0627-17-4) plates containing 50 ug/mL hygromycin (GoldBio, cat. #H-270) and incubated at 37° C for 3-5 days. Colonies were verified for insertion of the payload plasmid by PCR and subsequently confirmed by whole genome resequencing (SeqCenter).

For knockout strains, electrocompetent induced cells were co-transformed with pKM464 (Addgene # 108322)^37^ and a targeting oligo. The Δ*mceG* strain (bBEL594) harboring a deletion of *mceG* (*MSMEG_1366*) was generated using the following oligo (IDT Ultramer DNA Oligo): 5’-CCGTGACGGCCCGGCCTGATCGCGGCAAACTCACGCCTGCTTGGGCACCTCGATGACGCCGGTTTGTACCGTACACCACTGAGACCGCGGTGGTTGACCAGACAAACCCAACCCCGTCACGTCGATTTGGACGCCCATCAAAGATCCTTCCCGCTACGCCTACCACAC-3’.

Transformants were plated on 7H10 plates containing 50 ug/mL hygromycin and incubated at 37° C for 3-5 days. Colonies were verified for insertion of the payload plasmid by PCR and subsequently confirmed by whole genome resequencing (SeqCenter).

### Complementation plasmid construction

For complementation of the ORBIT-constructed *mceG* knockout (bBEL594), a derivative of pMV261zeo (a gift from Jeffory Cox at University of California, Berkeley) was cloned containing wild type *mceG* (pBEL2759). The coding sequence of *mceG* was amplified genomic DNA extracted from *Msmeg* cells using AccuPrime Pfx DNA Polymerase (Invitrogen, cat. #12344032) and cloned into pMV261zeo using Gibson assembly. TOP10 cells (Invitrogen, cat.# C404010) were transformed with the assembled vector using heat shock and plated on LB+agar plates containing 25 ug/mL zeocin (Gibco, cat. #R25001). Colonies were screened for correct DNA sequences using Sanger sequencing (Azenta). Complementation plasmids harboring MceG mutants were generated in a similar manner (pBEL2713, MceG(Y178A); pBEL2719, MceG(Δ242-360)).

Complementation plasmids were electroporated into electrocompetent Δ*mceG Msmeg* cells. Cells were plated on 7H10 plates containing appropriate antibiotics (e.g. 25 μg/mL zeocin, 50 μg/mL hygromycin). Colonies were selected, cultured in Middlebrook 7H9 (Difco, cat.#271310) containing 0.05% (v/v) Tween 80 (Sigma, cat. #P1754) and appropriate antibiotics, frozen as 20% glycerol stocks for future use.

### Cholesterol growth assay

Cholesterol growth assay was adapted from previous studies^14,35^. Briefly, *Msmeg* strains were streaked on 7H10 plates supplemented with 0.05% (v/v) Tween 80 and the appropriate antibiotics from frozen glycerol stocks. Colonies were used to seed M9 medium (1 L dH_2_O, 12.8 g Na_2_HPO_4_, 3 g KH_2_PO_4_, 0.5 g NaCl, 1 g NH4Cl, 25 μL 1 M CaCl_2_, 500 μL 1 M MgSO_4_) supplemented with 0.5% glycerol and 0.05% (v/v) tyloxapol (Ty, Sigma, cat. #T0307) with appropriate antibiotics. M9 cultures were grown to OD_600_ of ~0.7-1.0 at 37° C and harvested. Strains were washed twice by pelleting cells by centrifugation at 4,000 rcf for 5 mins at 22° C and resuspended in M9 medium with 0.05% tyloxapol. After the wash steps, strains were resuspended in M9 medium with 0.05% tyloxapol to an OD_600_ of 0.1 and were used to seed 200 μL cultures (starting OD_600_ of 0.005) for growth in 96-well plates. For each strain, the following medias were used: 1) M9+0.05% Ty+ 0.5% (v/v) glycerol (carbon source positive control), 2) M9+0.05% Ty+0.009 g/mL methyl-β-cyclodextrin (MBC, Sigma, cat. #C4555) (no carbon source control), and 3) M9+0.05% Ty+0.009 g/mL MBC+ 0.69 mM cholesterol (Sigma, cat. #C8667). Cultures were grown at 37° C with shaking and OD_600_ was monitored for each strain using a plate reader (BioTek). At least three biological replicates were conducted and plotted using Prism (GraphPad).

### Bacterial growth and protein purification

*Msmeg* was grown in Middlebrook 7H9 supplemented with 0.05% (v/v) Tween 80 and additional antibiotics as needed (e.g. 50 ug/mL hygromycin). For protein expression and purification of chromosomally GFP-tagged MceG (bBEL591) or GFP-tagged LucB (bBEL595), overnight cultures of each strain were diluted 1:1000 and grown with shaking at 37° C and 200 rpm until 0.8-1.2 OD_600_. Cells were harvested by centrifugation at 4,000 rcf, 4 °C. Pellets were resuspended in lysis buffer (50 mM Tris-HCl pH 7.5, 150 mM NaCl, 5 mM MgSO_4_, 5 mM 6-aminocaproic acid (Sigma, cat. #A2504), 5 mM benzamidine (Sigma, cat. #B6506) and 1 mM phenylmethylsulfonyl fluoride (PMSF, Sigma, cat. #10837091001)) and stored at −80 °C. Cells were thawed at room temperature and lysed by four passes through an chilled Emulsiflex-C3 cell disruptor (Avestin) at an output pressure of 20 kpsi. Unlysed cells and debris were removed by centrifugation at 39,000 rcf for 30 min at 4 °C. Membranes from the resulting supernatant were pelleted by ultracentrifugation in a Fiberlite F37L-8 x 100 Fixed-Angle Rotor (Thermo Scientific, cat. # 096-087056) at 37,000 rpm for 90 min at 4 °C and resuspended in membrane resuspension (MR) buffer (50 mM Tris-HCl pH 7.5, 15% (v/v) glycerol, 5 mM MgSO_4_, 150 mM NaCl, 5 mM 6-aminocaproic acid, 5 mM benzamidine, and 1 mM PMSF). Resuspended membranes were stored −80 °C. For affinity purification, membranes were thawed and solubilized overnight with addition of 20 mM n-dodecyl-β-D-maltoside (DDM, Inalco, cat. #D310S) at 4 °C and insoluble material was removed by centrifugation at 37,000 rpm for 60 min. GFP affinity resin was prepared using a method adopted from Pleiner et al.^74^. Briefly, purified His14-Avi-SUMO^Eu1^-anti GFP nanobody (expressed from pTP396, Addgene #149336)^74^ was biotinylated using BirA (expressed from pTP264, Addgene #149334)^74^ and further purified using a Superdex 200 16/60 gel filtration column (Cytiva, cat. # 28-9909-44) equilibrated in GF1 buffer containing: 50 mM Tris/HCl pH 7.5, 200 mM NaCl, 1 mM dithiothreitol (DTT, Amresco, cat. #M109). The biotinylated anti-GFP nanobody was added to Pierce High Capacity Streptavidin Agarose Resin (Thermo Scientific, cat. #20359) equilibrated in GF1 buffer and allowed to incubate with the resin overnight at 4 °C. 0.6 mL bed volume of resin was washed three times with GF1 buffer and blocked by incubation with 100 μM biotin (Sigma, cat. #B4501) in 50 mM HEPES/KOH pH 7.5 for 5 min on ice with occasional mixing. Beads were washed three times with GF1 Buffer and subsequently washed three times with MR buffer containing 20 mM DDM prior to use. Solubilized membranes were incubated with the equilibrated GFP affinity resin at 4 °C for 6 hours and then washed three times with 125 column volumes of membrane wash (MW) buffer (50 mM Tris-HCl pH 7.5, 15% (v/v) glycerol, 5 mM MgSO_4_, 150 mM NaCl, 5 mM 6-aminocaproic acid, 5 mM benzamidine, 1 mM DDM and 1 mM PMSF). Immobilized proteins were eluted by incubation with 1 mL of 250 nM SENP^EuB^ protease (expressed and purified from pAV286 (Addgene # 149333))^74^ overnight at 4° C. Eluted proteins were pooled and concentrated before separation on a Superdex 200 16/60 column (GE Healthcare) equilibrated in GF2 Buffer (50 mM Tris-HCl pH 7.5, 5 mM MgSO_4_, 150 mM NaCl, 1 mM DDM, and 1 mM DTT). Fractions containing GFP-tagged MceG or GFP-tagged LucB were buffered exchanged in storage buffer (50 mM Tris-HCl pH 7.5, 20% (v/v) glycerol 5 mM MgSO_4_, 150 mM NaCl, 1 mM DDM, and 1 mM DTT) and stored separately in −80 °C.

### Western blot for detection of GFP

Purified protein fractions were separated on a Mini-PROTEAN TGX Stain-Free protein gel (Bio-Rad Laboratories, Inc.). Separated protein bands were visualized using “Stain Free Gel” application mode on ChemiDoc MP Imaging System (Bio-Rad Laboratories, Inc.). Protein gel was transferred to a nitrocellulose membrane (Bio-Rad, cat. #1704271) using a Trans-Blot Turbo Transfer System (Bio-Rad Laboratories, Inc.). Membranes were blocked in PBST containing 5% milk for 30 min at 22 °C. The membranes were then incubated with primary antibodies for GFP (custom anti-GFP rabbit polyclonal (provided by Foley lab, Memorial Sloan Kettering Cancer Center) at a dilution of 1:5,000) and His (mouse anti-penta-His antibody (Qiagen, cat. #34660) at a dilution of 1:10,000) in PBST + 5% milk overnight at 4 °C. The membranes were washed three times with PBST and were incubated with goat antirabbit IgG polyclonal antibody (IRDye 800CW (LI-COR Biosciences cat. #925–32211) at dilution of 1:10,000) and goat anti-mouse IgG polyclonal antibody (IRDye 680RD, LI-COR Biosciences #926-68070 at a dilution of 1:10,000) as the secondary antibodies in PBST + 5% milk for 1 hr at 22° C. The membranes were washed three times with PBST and imaged using a LI-COR (LI-COR Biosciences) and analyzed by ImageJ ^75^.

### Negative stain electron microscopy

To prepare grids for negative stain electron microscopy, a fresh sample of either MceG-GFP or LucB-GFP was applied to a freshly glow discharged (30 seconds) carbon coated 400 mesh copper grid (Ted Pella Inc., cat. #01754-F) and blotted off. Immediately after blotting, a 2% uranyl formate solution was applied for staining and blotted off on filter paper. Application and blotting of stain was repeated five times. Samples were allowed to air dry before imaging. Data were collected on a Talos L120C TEM (FEI) equipped with a 4K x 4K OneView camera (Gatan) at a nominal magnification of 73,000x corresponding to a pixel size of 2.00 Å /pixel on the specimen, and a defocus range of −1 to −2 μm defocus. For LucB-GFP data, data processing was carried out in cryoSPARC v3.3.1^60^. Micrographs were imported, particles were picked manually as templates for Template Picking. Particles that were picked by template picking were sorted using 2D Classification.

### Sample preparation for mass spectrometry

Protein samples from wild-type *Msmeg* cells (strain mc^2^155, bBEL246), MceG-GFP strain (bBEL591), LucB-GFP (bBEL595) strain were purified using the protein purification method described above. Three biological replicates were performed for each strain and analyzed by mass spectrometry. Affinity purified proteins were reduced with DTT at 57 °C for 1 hour (2 μL of 0.2 M) and subsequently alkylated with iodoacetamide at room temperature in the dark for 45 minutes (2 μL of 0.5 M). To remove detergents and other buffer components the samples were loaded onto a NuPAGE® 4-12% Bis-Tris Gel 1.0 mm (Life Technologies Corporation). The gel was run for approximately 25 minutes at 200 V. The gel was stained using GelCode Blue Stain Reagent (Thermo Scientific). The entire protein band was excised, extracted and analyzed in a single mass spectrometry analysis per gel lane. The excised gel pieces were destained in 1:1 v/v solution of methanol and 100 mM ammonium bicarbonate solution using at least three exchanges of destaining solution. The destained gel pieces were partially dehydrated with an acetonitrile rinse and further dried in a SpeedVac concentrator until dry. 200 ng of sequencing grade modified trypsin (Promega) was added to each sample. After the trypsin was absorbed, 250 μL of 100 mM ammonium bicarbonate was added to cover the gel pieces. Digestion proceeded overnight on a shaker at room temperature. The solution was removed and placed into a separate Eppendorf tube. The gel pieces were covered with a solution of 5% formic acid and acetonitrile (1:2; v:v) and incubated with agitation for 15 min at 37°C. The extraction buffer was removed and placed into the Eppendorf tube with the previously removed solution. This was repeated three times and the solution dried in the SpeedVac concentrator. The samples were reconstituted in 0.5% acetic acid and loaded onto equilibrated Micro spin columns (Harvard apparatus) using a micro centrifuge. The bound peptides were washed three times with 0.1% TFA followed with one wash with 0.5% TFA. Peptides were eluted by the addition of 40% acetonitrile in 0.5% acetic acid followed by 80% acetonitrile in 0.5% acetic acid. The organic solvent was removed using a SpeedVac concentrator and the sample reconstituted in 0.5% acetic acid and kept at −80 °C until analysis.

### Mass spectrometry data collection

LC separation was performed online on an EASY-nLC 1200 (Thermo Scientific) utilizing Acclaim PepMap 100 (75 μm x 2 cm) precolumn and PepMap RSLC C18 (2 μm, 100A x 50 cm) analytical column. Peptides were gradient eluted directly to an Orbitrap Elite mass spectrometer (Thermo Fisher) using a 95 min acetonitrile gradient from 5 to 35 % B in 60 min followed by a ramp to 45% B in 10 min and 100% B in another 10 min with a hold at 100% B for 10 min (A=2% acetonitrile in 0.5% acetic acid; B=80% acetonitrile in 0.5% acetic acid). Flow rate was set to 200 nl/min. High resolution full MS spectra were acquired every three seconds with a resolution of 120,000, an AGC target of 4e5, with a maximum ion injection time of 50 ms, and scan range of 400 to 1500 m/z. Following each full MS data-dependent HCD MS/MS scans were acquired in the Orbitrap using a resolution of 30,000, an AGC target of 2e5, a maximum ion time of 200 ms, one microscan, 2 m/z isolation window, normalized collision energy (NCE) of 27, and dynamic exclusion of 30 seconds. Only ions with a charge state of 2-5 were allowed to trigger an MS2 scan.

### Analysis of mass spectrometry data

The MS/MS spectra were searched against the NCBI *Mycobacterium smegmatis* database with common lab contaminants and the sequence of the tagged bait proteins were added using SEQUEST within Proteome Discoverer 1.4 (Thermo Fisher). The search parameters were as follows: mass accuracy better than 10 ppm for MS1 and 0.02 Da for MS2, two missed cleavages, fixed modification carbamidomethyl on cysteine, variable modification of oxidation on methionine and deamidation on asparagine and glutamine. The data was filtered using a 1% FDR cut off for peptides and proteins against a decoy database and only proteins with at least 2 unique peptides were reported in Supplementary Table 2.

To obtain a probabilistic score (SAINT score) that a protein is an interactor of either MceG or LucB, the data were analyzed using the SAINT Express algorithm^59^. A one-sided volcano plot was generated showing fold change (Tag/WT) versus SAINT score. Proteins with a SAINT score ≥0.67 yielded an FDR of ≤5% and were considered potential interactors. Analyzed data are annotated in Supplementary Table 3 (for MceG) and in Supplementary Table 5 (for LucB) and plotted in Fig. 1f (for MceG) and Extended Data Fig. 9e (for LucB), respectively, using Prism (GraphPad).

### Cryo-EM sample preparation

The MceG-GFP complex was freshly purified as described above. Gel filtration fractions corresponding to higher-molecular weight complexes containing MceG were screened by negative-stain electron microscopy. Fractions of interest were then concentrated to ~1.7 mg/mL in cryo-EM buffer (50 mM Tris-HCl pH 7.5, 5 mM MgSO_4_, 150 mM NaCl, 1 mM DDM, and 1 mM DTT). Continuous carbon grids (Quantifoil R 2/2 on Cu 300 mesh grids + 2 nm Carbon, Quantifoil Micro Tools C2-C16nCu30-01) were glow-discharged for 5 sec in an easiGlow Glow Discharge Cleaning System (Ted Pella Inc.). 3.5 μL sample was added to the glow-discharged grid. Using a Vitrobot Mark IV (Thermo Fisher Scientific), grids were blotted for 3-3.5 seconds at 22 °C with 100% chamber humidity and plunge-frozen into liquid ethane. Grids were clipped for screening.

### Cryo-EM screening and data collection

Clipped cryo-EM grids were screened at NYU Cryo-EM Laboratory on a Talos Arctica (Thermo Fisher Scientific) equipped with a K3 camera (Gatan). Images of the grids were collected at a nominal magnification of 36,000x (corresponding to a pixel size of 1.0965 Å) with total dose of 50 e^-^ per Å^2^, over a defocus range of −2.0 to −3.0 μm. Grids were selected for data collection based on ice quality and particle distribution. Selected cryo-EM grids were imaged at Pacific Northwest Center for Cryo-EM on “Krios 2”, a Titan Krios G3 electron microscope (Thermo Fisher Scientific) equipped with a K3 BioContinuum direct electron detector (Gatan). Super-resolution movies were collected at 300 kV using SerialEM^76^ at a nominal magnification of 105,000x, corresponding to a super-resolution pixel size of 0.41275 Å (or a nominal pixel size of 0.8255 Å after binning by 2). Movies were collected over a defocus range of −0.8 to −2.4 μm and each movie consisted of 60 frames with a total dose of 60 e^-^ per Å^2^. A total of 43,925 movies were collected, consisting of 21,915 movies at 0° tilt and 22,010 movies at −30° tilt. Further data collection parameters are shown in Supplementary Table 4.

### Cryo-EM data processing

The dataset was split up into batches of 1,000 movies (45 batches total) and processed in cryoSPARC v3.3.1^60^, as described in figs. S3 and S4. Dose-fractionated movies were gain-normalized, drift-corrected, summed, and dose-weighted using the cryoSPARC Patch Motion module. The contrast transfer function was estimated for each summed image using cryoSPARC Patch CTF.

From the first batch of 1,000 images, 27 particles were manually picked in cryoSPARC that were then extracted (boxsize = 480 pixel (px)) and used to train within the Topaz Train module^77^ in cryoSPARC (expected number of particles = 50, use pretrained initialization, ResNet16). After training, particles were picked using the trained Topaz model and extracted (10,618 particles, box size = 480 px). CryoSPARC 2D classification (N = number of classes = 50) was performed and particles from 2D classes with high resolution detail were selected (1,051 particles) for Topaz Train (expected number of particles = 300, use pre-trained initialization, ResNet16). Trained Topaz model was used to pick and extract 105,604 (box size 480) particles that were curated by 2D classification (N = 50). Particles from the well-defined classes were selected (14,402 particles after removing duplicates) and further curated using 2D classification (N = 50).

Particles from classes representing top, side, and tilted views were selected (2,887 particles) and processed using cryoSPARC *Ab initio* Reconstruction to generate an initial 3D model (Ref 1: Complex (1,268 particles), Ref 2 (919 particles), Ref 3 (700 particles)). To generate decoys for downstream particle curation, 50,927 ‘junk’ particles were selected from the 2D classification and processed using cryoSPARC *Ab initio* Reconstruction to generate three decoy models (Decoy 1 (17,094 particles), Decoy 2 (16,915 particles), and Decoy 3 (16,918 particles)). For a more isotropic reconstruction in 3D, the 1,268 particles from Ref1 were sorted in 2D (N = 10) and different views of the particles were selected individually: side (588 particles), titled (505 particles), top (43 particles). These select particles were used to generate Topaz models to specifically pick side, tilted, and top views of the particle through the Topaz Train module (expected number of particles = 300, use pretrained initialization, ResNet16).

Using these Topaz picking models, separate Topaz Extract jobs were performed for each view, particles were extracted (box size 480, binned by 4), and combined. The combined particles were curated by cryoSPARC 2D classification (N = 50), subjected to duplicate removal (alignments2D), and curated in 3D using cryoSPARC Heterogenous Refinement (N = 4, templates = (1) Decoy1, (2) Decoy2, (3) Decoy3, (4) Model). Particles sorted into template 4 (Model) were selected for further processing. This curation scheme was performed on each batch of micrographs resulting in 2,869,223 curated particles, in which 1,820,584 particles came from the non-tilted images and the remaining 1,048,639 particles came from the −30° tilted images.

Particles were re-extracted (box size = 360 px, unbinned) and were further curated by running six rounds of Heterogeneous Refinement (N = 4, templates = (1) Decoy1, (2) Decoy2, (3) Decoy3, (4) Model), in which particles that were sorted into template 4 (Model) were used as input for the next round. After multiple rounds of Heterogeneous refinement (round 1: 992,273 particles, round 2: 637,446 particles, round 3: 510,255 particles, round 4: 468,001 particles, round 5: 437,324 particles, round 6: 414,343 particles) and removing remaining duplicates (alignment3D), the 341,566 curated particles were refined using cryoSPARC Non-Uniform Refinement^78^ generating a consensus map at 2.83 Å-resolution.

Heterogeneity was observed around the inner membrane (IM) region of the complex so particles were subject to a round of Heterogeneous Refinement (N = 4, templates = (1-4) consensus map). Class a (48,786 particles) and class b (113,261 particles) both contained additional density corresponding to extra protein density in the IM region and were combined, whereas the additional density were not observed in class c (59,724 particles) and class d (119,795 particles). Class c and Class d were very similar when compared by visual inspection, and these two classes were therefore combined. Non-uniform refinement was performed on the combined sets of particles, resulting in two major classes (both containing density for MceG, YrbE1AB, and Mce1ABCDEF): Class 1 that contains the extra protein density (162,047 particles, 2.94 Å) and Class 2 that lacks this density (179,519 particles, 3.04 Å).

Local refinements were performed for each class by recentering the particles on the region of interest using cryoSPARC Volume Alignment Tool, re-extracting the particles with the new center (box size = 360 px, unbinned), refining the particles on the re-centered 3D template using Non-uniform Refinement, performing particle subtraction in cryoSPARC using a mask around the region of interest, followed by refinement using cryoSPARC Local Refinement of the subtracted particles. This procedure was performed on each class to generate locally refined maps for the following regions: (i) MceG2, (ii) YrbE1AB+Mce1ABCDEF(transmembrane helix+transmembrane domains+Mce ring)+/-extra factor, (iii) Mce1ABCDEF(Mce ring+ first half of C-terminal Mce needle), (iv) Mce1ABCDEF (second half of C-terminal Mce needle). For class 1, the following maps were generated for corresponding regions: (i) Map1a (161,434 particles, 3.05 Å), (ii) Map1b (162,004 particles, 2.89 Å), (iii) Map1c (158,508 particles, 2.97 Å), (iv) Map1d (156,741 particles, 3.16 Å). For Class 2, the following maps were generated for each region: (i) Map2a (178,844 particles, 3.13 Å), (ii) Map2b (179,480 particles, 2.99 Å), (iii) Map2c (175,490 particles, 3.06 Å), (iv) Map2d (173,315 particles, 3.19 Å). To generate a composite map, particles from each class were re-extracted with a box size of 640 px (unbinned) and refined using Non-Uniform Refinement to generate maps that included the entire complex (Map1e for Class 1 and Map2e for Class 2). These maps were used as a template to stitch the locally refined maps together to generate a composite density map. In regions aside from the extra density (later assigned as LucB/MSMEG_3032), these maps were lower resolution compared to the map from the consensus set of particles before classification, but did not show any notable differences compared with the consensus map. Therefore, local refinements were performed on the consensus set of particles in similar fashion used to generate maps for model building, but with masking out the MSMEG_3032/LucB density.

Local refinements were performed using the same approach that was applied to Class 1 and Class 2 on the set particles from the consensus refinement. This procedure was utilized on the following regions: (i) MceG2, (ii) YrbE1AB+Mce1ABCDEF(transmembrane helix+transmembrane domains+Mce ring) masking out density for the extra factor, (iii) Mce1ABCDEF(Mce ring+ first half of C-terminal Mce needle), (iv) Mce1ABCDEF (second half of C-terminal Mce needle). For the consensus map, the following locally refined maps were generated for each region: (i) Map0a (340,238 particles, 2.91 Å), (ii) Map0b (341,490 particles, 2.73 Å), (iii) Map0c (332,050 particles, 2.75 Å), (iv) Map0d (330,104 particles, 3.00 Å). To generate a composite map, the consensus set of particles were also re-extracted with a box size of 640 px (unbinned) and refined using Non-Uniform Refinement to generate a map that included the entire complex (Map0e). This map was used as a template to stitch the locally refined maps together to generate a composite density map. These maps were of much higher quality compared to local refined maps of class 1 and class 2, thus used for initial model building.

For each map, the overall resolution reported in cryoSPARC was estimated using the gold-standard Fourier Shell Correlation criterion (FSC = 0.143). Directional FSCs were computed using 3DFSC^65^. Local resolution maps were computed using the cryoSPARC Local Resolution Estimation module. Locally refined maps were combined into composite maps for the consensus map, Class 1 and Class 2 using PHENIX v1.20.1 ‘Combine Focused Maps’ module^64^. Composite maps were generated for sharpened maps and half maps (for calculating FSC and estimating local resolution of the composite maps). For the consensus composite map, maps 0a, 0b, 0c, and 0d were combined using Map0e as a template to generate Map0. For the class 1 composite map, maps 1a, 1b, 1c, and 1d were combined using Map1e as a template to generate Map1. For the class 2 composite map, maps 3a, 3b, 3c, and 3d were combined using Map2e as a template to generate Map2. Global FSCs were calculated by importing composite half maps into the ‘Validation FSC’ cryoSPARC module and local resolution was estimated using the ‘Local Resolution’ cryoSPARC module. The nominal global resolution was estimated to be 2.71 Å for Map0, 2.76 Å for Map1 and 2.90 Å for Map2. Directional FSCs for the composite maps were computed using 3DFSC in cryoSPARC.

### Model building and refinement

The mass spectrometry data indicated a mixture of Mce1 and Mce4 proteins in the cryo-EM sample. To assess which proteins were present in the cryo-EM reconstruction, their stoichiometry and position in the complex, we generated AlphaFold2^63^ predictions for each MCE-related protein and assessed their fit into the consensus reconstruction, which contains the ATP-binding cassette (ABC) transporter and the MCE ring. Using ColabFold^79^, AlphaFold ^63^ predictions were generated for MceG (AFpdb1), Mce1 proteins (AFpdb1-9), Mce4 proteins (AFpdb10-17), and orphaned MCE protein (AFpdb18). Predictions are summarized in Supplementary Table 6. We performed rigid-body fits of the predicted structures into the cryo-EM map using UCSF Chimera v1.16^80^, and determined that the complex consisted of two protomers of MceG, two protomers of YrbEs, and six MCE proteins. The two protomers of MceG (AFpbd1) fit unambiguously into the density that corresponded to the ATPase component of the ABC transporter. For YrbE and MCE proteins, we further refined the rigid-body fitted models using real-space refinement in PHENIX v1.20.1^64^. We then examined regions of each protein where the sequences are divergent between candidate proteins and used side chain density in order to assign the correct subunit. The YrbE subunits (AFpdb2-3,10-11) were fit as rigid bodies into the transmembrane region of the cryo-EM map using UCSF Chimera and refined in real space using PHENIX. The refined models were manually inspected in COOT v0.8.9.2^81^ to assess the overall fit for the Ca backbone and side chains of each protein into the map. Based on manual inspection, we assigned the cryo-EM density to YrbE1A and YrbE1B. The MCE domains of each Mce1 (AFpdb4-9) or Mce4 (AFpdb12-17) protein were fitted into each position of the MCE ring (positions 1-6) using UCSF Chimera. Once fit into the density, the MCE domains were real-space refined in PHENIX and manually inspected in COOT. Based on this analysis, Mce1 proteins fit best into the map and were assigned the following positions in the MCE ring (going clockwise): 1) Mce1A, 2) Mce1E, 3) Mce1B, 4) Mce1C, 5) Mce1D, 6) Mce1F. Thus, using this approach, we are able to unambiguously assign Mce1 protein subunits into the cryo-EM map (Extended Data Fig. 4a). Notably, oMce1A (AFpdb18), which was identified in the mass spectrometry data and is 84% identical to Mce1A, fits well into the cryo-EM map at the same position as Mce1A, suggesting a possible mixture of Mce1A and oMce1A in the reconstruction. Focused 3D classification around regions that differ between the two proteins did not produce classes where the density was resolved enough to unambiguously assign Mce1A versus oMce1A. Mce1A was used for modeling the Mce1 complex since it belongs in the same operon as the other Mce1 proteins.

As a starting point for model building of the entire complex, AlphaFold2^63^ and AlphaFold-Multimer^82^ were used to predict 3D structures of Mce1 proteins and subcomplexes as summarized in Supplementary Table 6. Predictions were performed on ColabFold^79^ and COSMIC^2 83^. The C-terminal region of AFpdb20 was trimmed starting at the following residues: Mce1A (residue 167), Mce1B (residue 151), Mce1C (residue 149), Mce1D (residue 160), Mce1E (residue 169), Mce1F (residue 149). For initial model building, AFpdb19, AFpdb20 (trimmed) and AFpdb21 were stitched together in PyMOL Molecular Graphics System (version 2.5.1 Schröodinger, LLC). Briefly, chains were renamed for each prediction: Mce1A (chain A), Mce1B (chain B), Mce1C (chain C), Mce1D (chain D), Mce1E (chain E), Mce1F (chain F), MceG (chain G and H), YrbE1A (chain I), YrbE1B (chain J). Predicted models were aligned in PyMOL using the ‘align’ command: 1) AFpdb19 and AFpdb20 were aligned based on chain I and J, and 2) AFpdb3 was aligned to AFpdb2 based the first α-helical module of the MCE proteins (chain A 150-167, chain B 134-151, chain C 134-149, chain D 145-160, chain E 151-169, chain F 135-149). Overlapping residues were trimmed and aligned models were stitched to produce a composite PDB of the Mce1 complex based on AlphaFold2 predictions.

From the three cryo-EM maps (Map0, Map1, Map2), Map0 has the highest resolution and most featureful density. Thus, modeling of the Mce1 complex was performed on the locally refined maps corresponding to Map0 (Map0a-d), except for model building of LucB, which was carried out using Map1b. Note that Map0 includes Mce1 complex particles with and without LucB. However, since there is no conformational change in the Mce1 complex at the resolutions we are at, the higher number of particles results in better quality density for the Mce1 complex minus LucB. Starting models were fitted into their corresponding locally-refined maps using the “Fit in Map” function in UCSF Chimera. For each map, the PDB was trimmed to remove regions of the protein that were not defined in the map. Rigid-body fitting into the cryo-EM maps was performed using PHENIX. Fitted models were visually inspected and manually adjusted in COOT. Real-space refinement with Ramachandran and secondary structure restraints was carried out in PHENIX using 5 cycles and 100 iterations to optimize the fit and reduce clashes. These models were iteratively inspected, manually rebuilt in COOT and refined in PHENIX until completion. Models built into the locally refined maps were aligned and stitched together in PyMOL. These models served as templates to generate a composite density map (Map0) for the consensus set of particles using the PHENIX ‘Combine Focused Maps’ module.

In Map0, poly-carbon chain unknown ligands (UNLs) were manually built into extra densities corresponding to substrates, and real-space refined in COOT. Elongated ligands (LIG, Chemical string: CCCCCCCCCCCCCCCCCCCCCCCCCCCC CCCCCCCCCCCC) were generated using PHENIX eLBOW^84^. Planar ligands derived from BNZ (benzene) and DKM (5-[(3S,4S)-3-(dimethylamino)-4-hydroxypyrrolidin-1-yl]-6-fluoro-4-methyl-8-oxo-3,4-dihydro-8H-1-thia-4,9b-diazacyclopenta[cd]phenalene-9-carboxylic acid). The composite model (containing ligands) was real-space refined into Map0 using PHENIX with global minimization, Ramachandran, secondary structure, and ligand restraints. We use UNLs because the resolution of our density clearly indicates the presence of additional molecules, but is not high enough to unambiguously define these molecules.

Our final consensus model for Map0 is nearly complete, apart from regions in Mce1A (residues 1-17), Mce1C (residues 310-524), Mce1D (residues 1-41 and 268-547), Mce1E (residues 1-32), Mce1F (residues 400-518), MceG protomers (residues 1, 256-280, and 326-360), YrbE1A (residues 1-13), and YrbE1B (residues 1-26), which are predicted to be flexible or unstructured (Extended Data Fig. 4d). Notably, no transmembrane helix was observed for Mce1E (MSMEG_0138; Rv0173/LprK in *Mtb*). Mce1E has been proposed to be a lipoprotein due the presence of a possible signal peptide and lipobox at its N-terminus^85^. Intriguingly, the first resolvable residue for Mce1E is C33, the cysteine that would be lipidated; however, density around this region was not sufficient to resolve this modification. In our mass spectrometry data, we do not detect N-terminal peptides for Mce1E which suggest that this region may indeed be cleaved.

Models for Map1 and Map2 were built using the model for Map0 as the starting model. The Map0 model was fitted and trimmed into the locally refined maps of each class in UCSF Chimera and PyMOL. Real-space refinement with Ramachandran and secondary structure restraints was carried out in PHENIX. Models were iteratively inspected, manually rebuilt in COOT, and refined in PHENIX until completion. For Class 1, extra protein density was observed near the TM of Mce1C in the inner membrane region of Map1b that corresponded to an additional subunit bound to the complex, LucB. To determine the identity of this unknown protein, we used a combination of model building and AlphaFold2. The Cα backbone of the polypeptide was traced manually in COOT. This Cα model was used to search structural databases (AlphaFold/Swiss-Prot v2, AlphaFold/Proteome v2, PDB100 211201, GMGCL 2204) using TM-align mode in Foldseek^57^. One of the highest-ranking hits from this search (TM-score 0.9509) was a putative, converserved, integral membrane protein from *Mycobacterium tuberculosis* (Rv2536, AF-P95017-F1-model_v2.pdb) found from the AlphaFold Protein Structure Database. The structure of the *Msmeg* ortholog of this protein (MSMEG_3032/LucB, AFpdb22) was predicted in ColabFold, docked into the cryo-EM density using Chimera, stitched into the model of Map1 using PyMOL), and refined in PHENIX. Completed locally refined models were then aligned and stitched together in PyMOL and used to generate a composite density map for Class 1 (Map1) and Class 2 (Map2) in PHENIX. Ligands were added to stitched models for Map1 and Map2 and models were real-space refined using PHENIX.

Statistics for the final models (Supplementary Table 4) were extracted from the results of the real_space_refine algorithm in PHENIX^64^ as well as MolProbity^86^ and EMringer^87^. Structural alignments and associated RMSD values were calculated using UCSF Chimera v1.16^80^ and PyMOL (Schröodinger, LLC). FSCs that were calculated in cryoSPARC were plotted in GraphPad Prism v9.3.1. Mce1 tunnel volume was calculated using CASTp v3.0^61^ with a probe radius of 2.2 Å and the inner diameter was calculated using MOLE v2.5 “pore mode”^68^. Cavity of the ABC transporter substrate-binding pocket calculated by CASTp v3.0 using a probe radius of 2.2 Å. Figures and Supplementary Videos were generated with PyMOL (Schröodinger, LLC), UCSF Chimera and ChimeraX^62^.

### Figure preparation

Figures in which map density is shown were prepared using ChimeraX^62^ with the following parameters:

- Fig. 2f. Map0 rendered with contour level 10.0.
- Fig. 3c. Ligand density from Map0 rendered using ChimeraX ‘volume zone’ with 3.0 Å distance cutoff around UNL1 and 7.6 contour level.
- Fig. 3f. Ligand density from Map0 was rendered using ChimeraX ‘volume zone’ with 3.0 Å distance cutoff around UNL1-31 and 7.0 contour level.
- Fig. 4c. Ligand density from Map0 rendered using ChimeraX ‘volume zone’ with 2.5 Å distance cutoff around UNL9 and 5.0 contour level.
- Fig. 5a. Map1 rendered with contour level 10.0. Map2 rendered with contour level 10.0.
- Fig. 5b. Protein density from Map1 rendered using ChimeraX ‘volume zone’ with 2.5 Å distance cutoff around 3D model of poly-alanine Cα backbone and 8.0 contour level.
- Extended Data Fig. 3a. Locally refined maps for the consensus set of particles were contoured with the following levels: Map0a (0.281), Map0b (0.257), Map0c (0.259), Map0d (0.199), Map0e (0.17).
- Extended Data Fig. 3b. Map0 contoured to 12.7.
- Extended Data Fig. 3e. Locally refined maps for Class 1 were contoured with the following levels: Map1a (0.172), Map1b (0.201), Map1c (0.185), Map1d (0.167), Map1e (0.15).
- Extended Data Fig. 3f. Map1 contoured to 10.1.
- Extended Data Fig. 3i. Locally refined maps for Class 2 were contoured with the following levels: Map2a (0.177), Map2b (0.148), Map2c (0.163), Map2d (0.126), Map2e (0.15).
- Extended Data Fig. 3j. Map2 contoured to 10.2.
- Extended Data Fig. 4a. Protein densities rendered using ChimeraX ‘volume zone’ with 2.0 Å distance cutoff around the indicated protein residues with the following contour levels: Mce1A/oMce1A (6.0), Mce1F (14.0), Mce1E (10.0), MceG protomer 2 (10.0), Mce1C (8.0), MceG protomer 1. YrbE1A (12.0), Mce1D (8.0), Mce1B (8.0), YrbE1B (10.0).
- Extended Data Fig. 4b. Ligand densities rendered using ChimeraX ‘volume zone’ with 2.5 Å distance cutoff around UNLs and with the following contour levels: UNL1 (8.0), UNL4 (6.0), UNL20 (8.0).
- Extended Data Fig. 4c. Protein densities rendered using ChimeraX ‘volume zone’ with 2.5 Å distance cutoff around each TM LucB and contour level 7.0.
- Extended Data Fig. 4d. Map0 contoured to 10.0.
- Extended Data Fig. 7c. Protein densities rendered using ChimeraX ‘volume zone’ with 2.0 Å distance cutoff around each PLL at contour level 10.0.
- Extended Data Fig. 8d. Protein densities rendered using ChimeraX ‘volume zone’ with 2.0 Å distance cutoff around YrbE1B C-terminus and Mce1F PLL and 8.7 contour level.

### Quantification and Statistical Analysis

The local resolution of the cryo-EM maps was estimated using cryoSPARC Local Resolution^60^. Directional 3DFSCs were calculated using 3DFSC^65^. The quantification and statistical analyses for model refinement and validation on deposited models were performed using PHENIX^64^, MolProbity^86^, and EMRinger^87^. Structural alignments and associated RMSD values were calculated using UCSF Chimera^80^ and PyMOL (Schröodinger, LLC). Tunnel and cavity volumes were calculated using CASTp v3.0^61^ and tunnel diameter was estimated using MOLE v2.5^68^. Multiple sequence alignments were generated using MUSCLE^69^ and JalView^70^. Phenotypic assays were replicated at least three times (*n* = 3). The mean and standard error of three replicates were plotted using Prism (GraphPad). Protein pulldowns were replicated at least three times (*n* = 3). MS data was analyzed using Proteome Discoverer 1.4 (Thermo Fisher Scientific) and SAINT Express algorithm^59^ and plotted using Prism (GraphPad).

## Supporting information

PDB 8FEF

PDB 8FED

PDB 8FEE

## Acknowledgements

We thank members of the Bhabha/Ekiert labs for helpful discussions and Nicolas Coudray, Juliana Ilmain, Georgia Isom, Mark Macrae, Fred Rubino, and Joe Sudar for feedback on our manuscript. We thank Heran Darwin (NYU School of Medicine) for supplying the *Mycobacterium smegmatis* strain (mc^2^155), Jeffery Cox (University of California, Berkeley) and Casey Vieni (NYU School of Medicine) for sharing plasmids, and the Foley Lab (Memorial Sloan Kettering Cancer Center) for providing purified rabbit GFP antibody. We thank Kristen Dancel-Manning for the illustration of the mycobacterial cell envelope; Fengxia Lang and Kristen Dancel-Manning from the NYU Microscopy Laboratory for overseeing use of the Talos L120C microscope and the facility in which the Talos L120C is housed; Alice Paquette, Willian Rice, and Bing Wang from the NYU Cryo-EM Laboratory for assistance with cryo-EM grid screening and microscope operation; Sean Mulligan and Lauren Vega at the Pacific Northwest Center for Cryo-EM for assistance with cryo-EM data collection. EM data processing has utilized computing resources at the HPC Facility at NYU, and we thank the HPC team for high-performance computing support. We thank the Central Lab Services team at NYU School of Medicine for preparation of media and buffers.

This work was supported by the following funding sources: PEW-00033055 (NIH/NIGMS, to G.B.), Schmidt Science Fellows (to J.C.) and pilot funding from the NYU Langone Health Antimicrobial-resistant Pathogen Program (to G.B. and D.C.E.). The NYU Microscopy Center is partially supported by NYU Cancer Center Support Grant NIH/NCI P30CA016087. The mass spectrometric experiments were supported in part by NYU Grossman School of Medicine and with a shared instrumentation grant from the NIH, 1S10OD010582-01A1 for the purchase of an Orbitrap Fusion Lumos. A portion of this research was supported by NIH grant U24GM129547 and performed at the PNCC at OHSU and accessed through EMSL (grid.436923.9), a DOE Office of Science User Facility sponsored by the Office of Biological and Environmental Research

## Author contributions

J.C., D.C.E., and G.B. conceived the project. D.C.E. and G.B. supervised and administered project. J.C. performed cloning, protein purifications and biochemistry. J.C. prepared cryo-EM specimens, collected and processed cryo-EM data. J.C., D.C.E., G.B. built models and performed structural analysis. J.C., A.F. and C.F performed phenotypic assays. J.P. and B.U. carried out mass spectrometry experiments and analyses. J.C., B.U., D.C.E., G.B. acquired funding for the project. J.C., D.C.E., G.B. wrote the original draft of the manuscript. J.C., A.F., C.F, B.U., D.C.E., G.B. revised and edited manuscript.

## Supplementary information

Supplementary Information is available for this paper.

## Data and materials availability

Correspondence and requests for materials should be addressed to Damian C. Ekiert and Gira Bhabha. The cryo-EM maps have been deposited in the Electron Microscopy Data Bank with accession codes: Map0 (EMD-29025), Map0a (EMD-29228), Map0b (EMD-29229), Map0c (EMD-29230), Map0d (EMD-29231), Map0e (EMD-29232), Map1 (EMD-29023), Map1a (EMD-29233), Map1b (EMD-29234), Map1c (EMD-29235), Map1d (EMD-29236), Map1e (EMD-29237), Map2 (EMD-29024), Map2a (EMD-29238), Map2b (EMD-29239), Map2c (EMD-29240), Map2d (EMD-29241), and Map2e (EMD-29242). The coordinates of the atomic models have been deposited in the Protein Data Bank under accession codes: PDB 8FEF (model for Map0), PDB 8FED (model for Map1), PDB 8FEE (model for Map2). Cryo-EM data was deposited in Electron Microscopy Public Image Archive: EMPIAR-11343. The mass spectrometry files are available at MassIVE (https://massive.ucsd.edu) with dataset identifier MSV000090807 and ProteomeXchange (proteomexchange.org) with identifier PXD038456. Bacterial strains and plasmids have been deposited in Addgene and identifiers are listed in Supplementary Table 1.

