## Extended Data Figures for "Structure of an endogenous mycobacterial MCE lipid transporter"

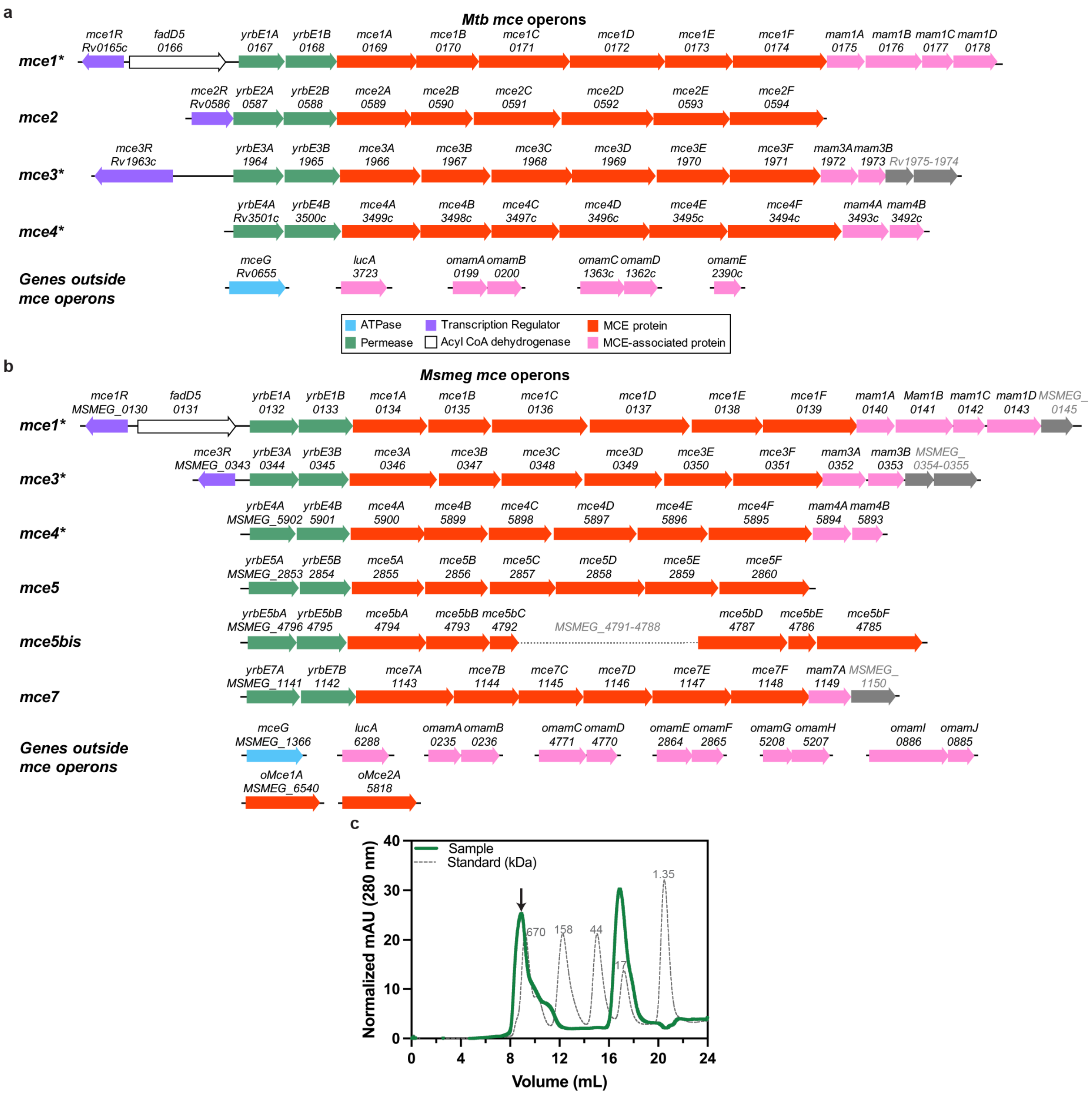


**Extended Data Fig. 1: MCE systems in *Mycobacterium tuberculosis and Mycobacterium smegmatis.*** **a**, Schematic of the four Mammalian Cell Entry (MCE) operons in the *Mycobacterium tuberculosis* (*Mtb)* genome. Arrows represent individual genes and their direction. Genes colored as indicated the key below. Operons annotated with an asterisk are conserved between *Mtb* and *Mycobacterium smegmatis* (*Msmeg)*. **B**, Schematic of the six MCE operons in the *Msmeg* genome. Genes colored as in Extended Data Fig. 1a. **c**, Size exclusion chromatogram of MceG-GFP pulldown (green) superimposed with protein standards (grey dotted line). Black arrow indicates protein sample shown in Fig. 1e and analyzed by mass spectrometry in Fig. 1f.

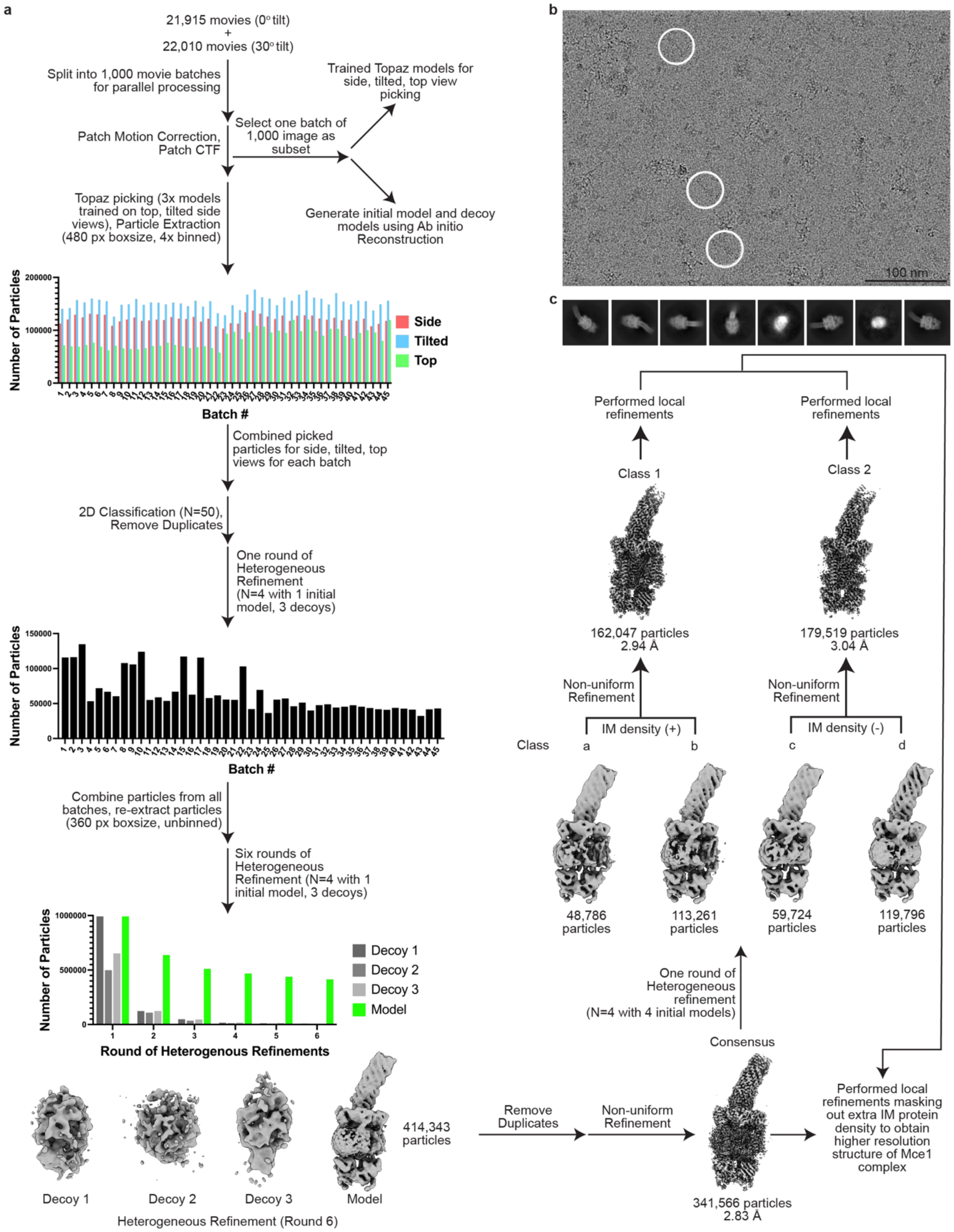


**Extended Data Fig. 2: Cryo-EM data processing workflow (Part 1). a**, Cryo-EM data processing pipeline. **b**, Representative cryo-EM micrograph. Particles of interest are circled in white. Scalebar (100 nm) is indicated on the bottom right of the micrograph. **c**, Representative 2D classes of complex. Eight 2D class averages showing different views of the particles were generated in cryoSPARC[60] using the final set of 'consensus' particles extracted with box size of 360 pixels and no binning.

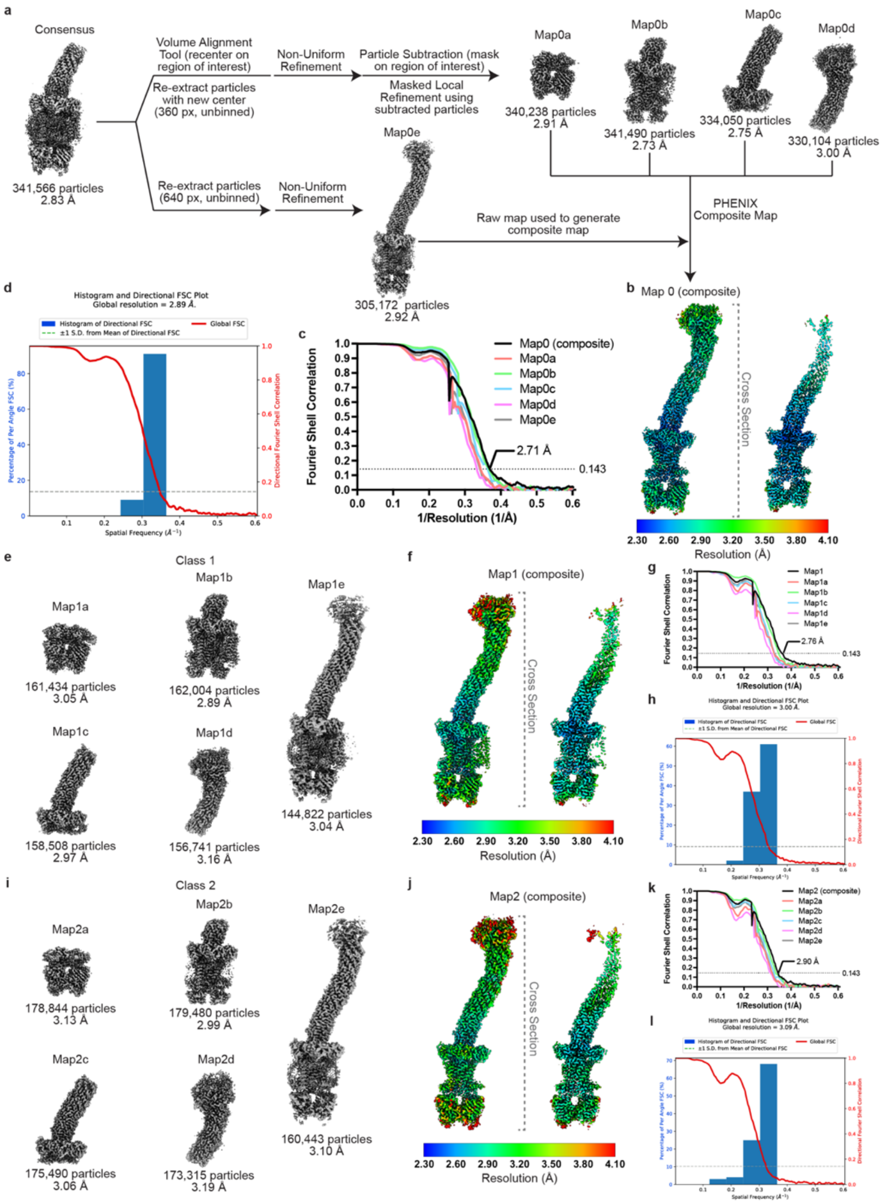


**Extended Data Fig. 3: Cryo-EM data processing workflow (Part 2). a**, Processing pipeline for local refinements performed in cryoSPARC[60] and composite map generation. Locally refined maps and raw map for consensus set of particles are shown. Map labels are indicated above the map and particle count and average resolution as reported by cryoSPARC are shown below. Composite density maps were generated in PHENIX[64]. **b**, Composite Map0 colored by local resolution that was estimated in cryoSPARC. (left) Whole structure view. (right) Cross-sectional view. (bottom) key for local resolution coloring, ranging from 2.30 Å (blue) to 4.10 Å (red). **c**, Gold-standard FSC curve calculated in cryoSPARC for composite, locally refined, and raw maps for consensus set of particles. The dotted line represents the 0.143 FSC cutoff. **d**, Directional 3DFSC[65] calculated for Map0 (composite). **e**, Locally refined and raw maps for Class 1. Map labels are indicated above the map and particle count and average resolution as reported by cryoSPARC are shown below. **f**, Composite density map for Class 1 (Map1) colored by local resolution that was estimated in cryoSPARC. (left) Whole structure view. (right) Cross-sectional view. (bottom) key for local resolution coloring, ranging from 2.30 Å (blue) to 4.10 Å (red). **g**, Gold-standard FSC curve calculated in cryoSPARC for composite, locally refined, and raw maps for Class 1. The dotted line represents the 0.143 FSC cutoff. **h**, Directional 3DFSC calculated for Map1 (composite). **i**, Locally refined and raw maps for Class 2. Map label is indicated above the map and particle count and resolution are shown below. **j**, Composite density map for Class 2 (Map2) colored by local resolution that was estimated in cryoSPARC. (left) Whole structure view. (right) Cross-sectional view. (bottom) key for local resolution coloring, ranging from 2.30 Å (blue) to 4.10 Å (red). **k**, Gold-standard FSC curve calculated in cryoSPARC for composite, locally refined, and raw maps for Class 2. The dotted line represents the 0.143 FSC cutoff. **l**, Directional 3DFSC calculated for Map2 (composite).

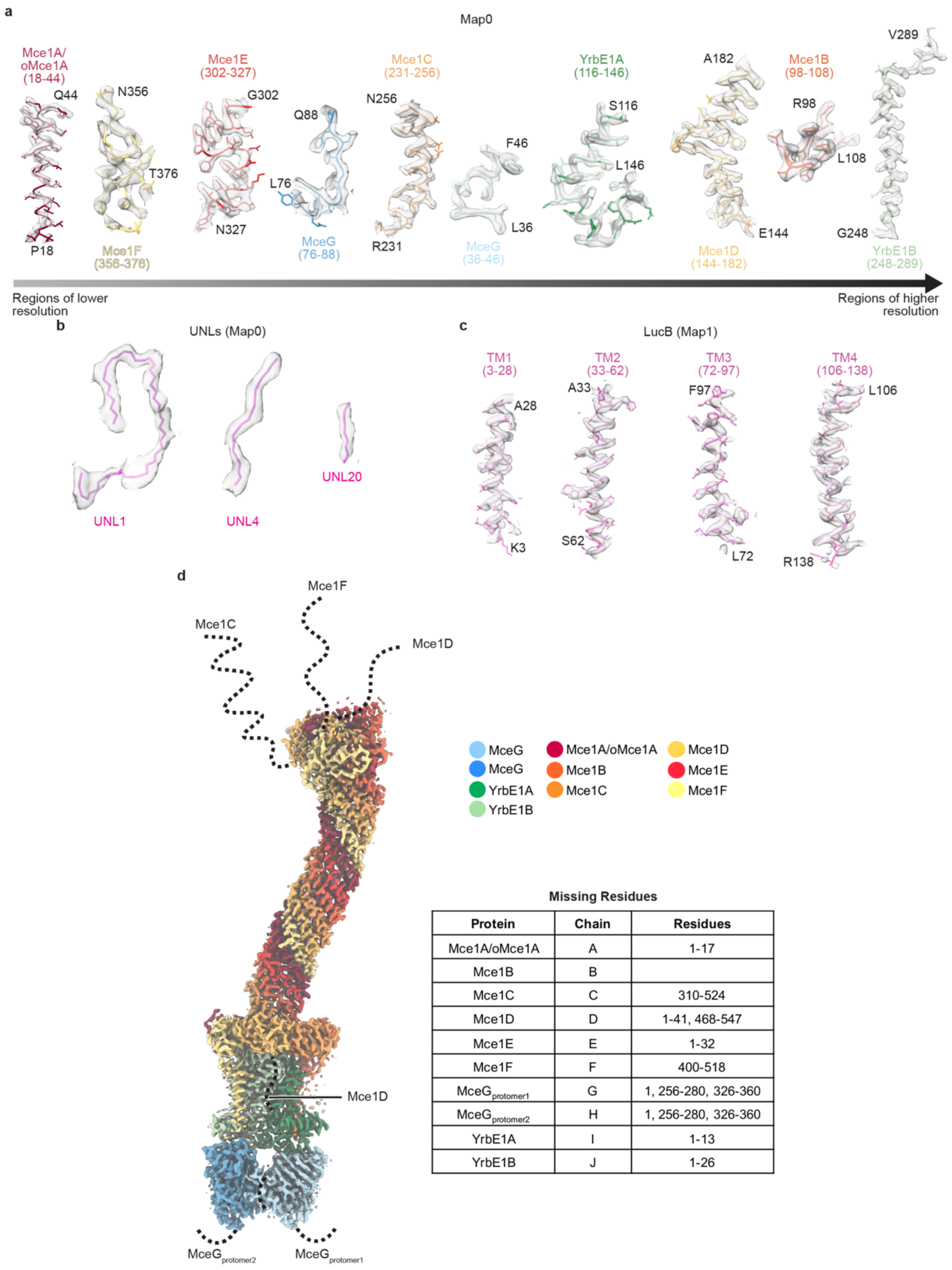


| Protein | Chain | Residues |
|---|---|---|
| Mce1A/oMce1A | A | 1-17 |
| Mce1B | B | |
| Mce1C | C | 310-524 |
| Mce1D | D | 1-41, 468-547 |
| Mce1E | E | 1-32 |
| Mce1F | F | 400-518 |
| $MceG_{protomer1}$ | G | 1, 256-280, 326-360 |
| $MceG_{protomer2}$ | H | 1, 256-280, 326-360 |
| YrbE1A | I | 1-13 |
| YrbE1B | J | 1-26 |

**Extended Data Fig. 4: Representative density from areas at different resolutions and missing regions in our maps. a**, Examples are shown from each subunit for Map0. Examples are arranged left to right from lower to higher resolution. **b**, Examples of ligand density from Map0. **c**, Examples of fits for LucB transmembrane helices (TM) from Map1. **d**, Missing regions in consensus composite density map (Map0). Map0 color-coded based on the key on the right. Regions of the proteins that were not resolvable in the cryo-EM map are indicated by dotted lines, drawn approximately to scale, and indicated in the chart.

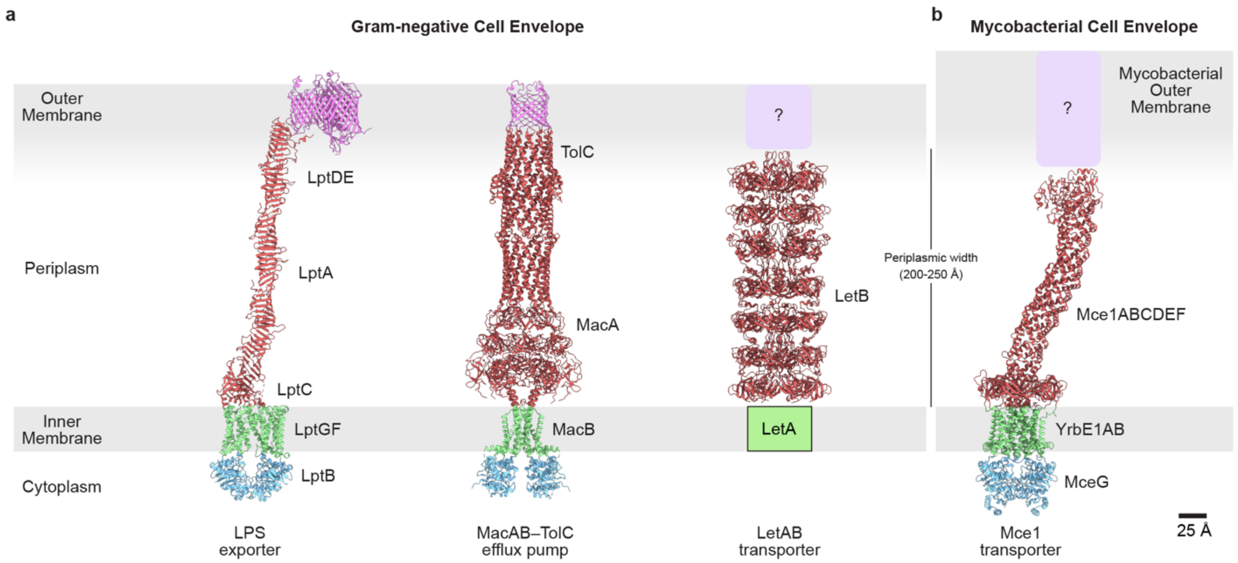


**Extended Data Fig. 5: Comparison of periplasm-spanning transporters in double-membraned bacteria. a**, Protein complexes that span the cell envelope in Gram-negative bacteria. (left) LPS exporter (modeled based on PDBs 6MIT[23], 2R19[66], 5IV9[67]); (middle) MacAB-TolC efflux pump (PDB 5NIK[27]); (right) LetAB transporter (PDB 6V0D[43]). **b,** Structure of mycobacterial Mce1 transporter.

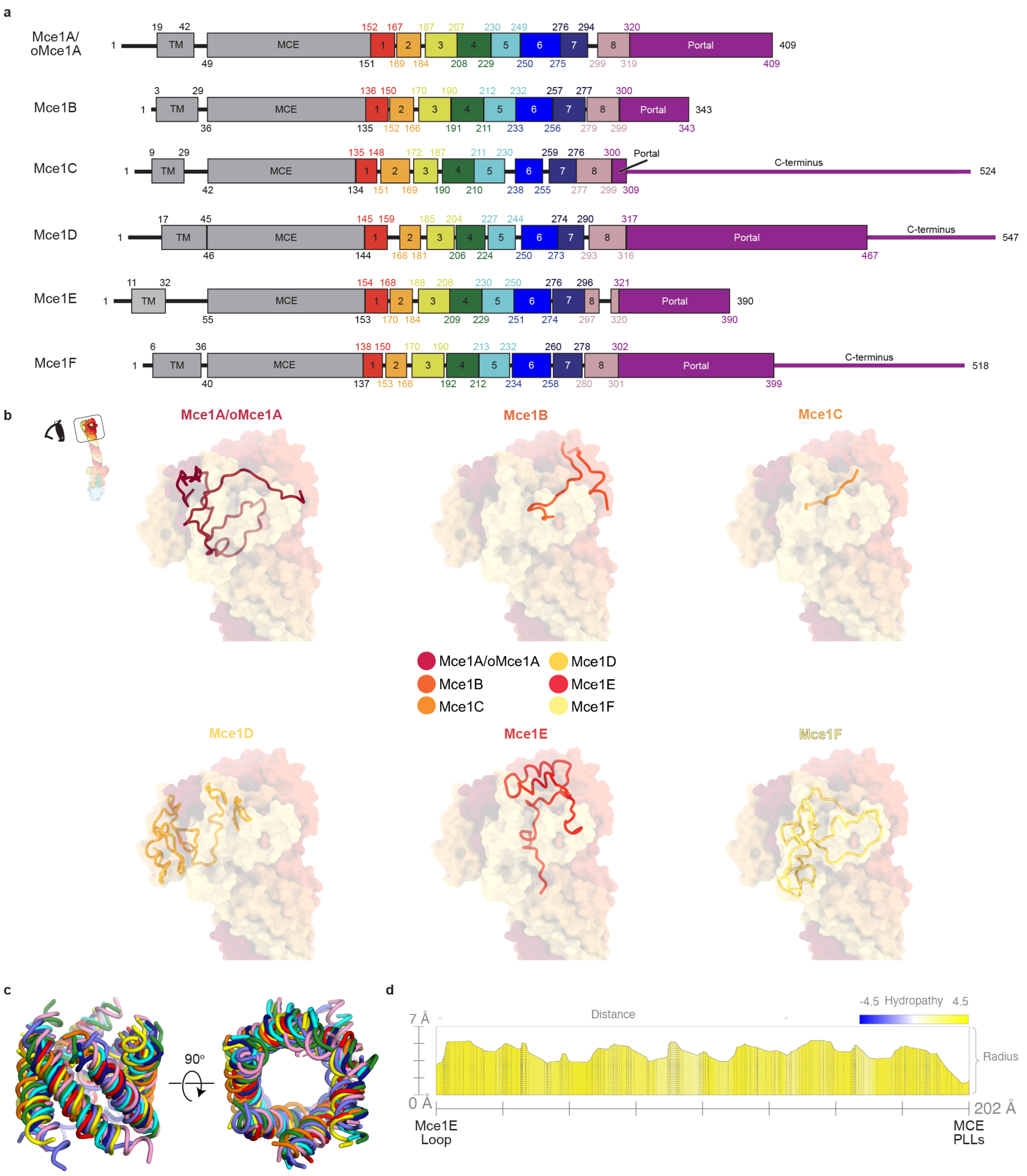


**Extended Data Fig. 6: Structural features of Mce1ABCDEF portal and needle modules in the Mce1 complex. a**, Schematic of Mce1ABCDEF proteins showing boundaries for the transmembrane helix (TM), MCE domain (MCE), eight needle modules, and portal domain/C-terminus. **b**, C-termini of Mce1ABCDEF that form the portal domain as indicated in upper-left inset. Proteins are shown as transparent molecular surfaces and colored as in the key. Superimposed in each panel are the residues of the indicated protein, shown as cartoon. **c**, Structural alignment of the eight structural modules in PyMOL using 'cealign' command (Schrödinger, LLC.). Proteins are superimposed, and colored as in Extended Data Fig. 6a. **d**, Hydrophobicity and radius of tunnel in Mce1 needle calculated using MOLE 2.5[68]. X-axis shows distance going down the tunnel (from left to right); Y-axis shows the distance away from the central axis of the tunnel. Plot shows hydrophobicity, where blue is hydrophilic and yellow is hydrophobic.

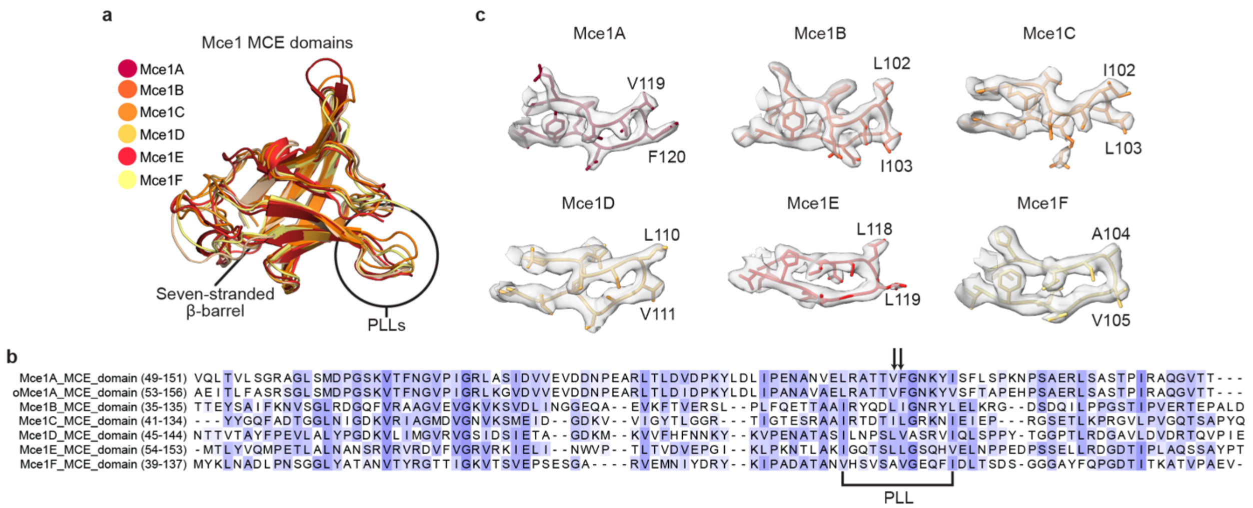


**Extended Data Fig. 7: Structural features of MCE domains. a**, Structural alignment of Mce1 MCE domains color-coded based on key. Domains were aligned to the MCE domain of Mce1A. Pore-lining loops (PLLs) are circled. **b**, Protein sequence alignment of the MCE domains from *Msmeg* Mce1 proteins using MUSCLE[69] and visualized using JalView[70]. Sequence alignment is colored by BLOSUM62 score (not conserved, white; conserved, blue). PLL region is highlighted with black bracket and pore facing residues are indicated with black arrows. **c**, Gallery of Mce1 PPLs. Protein backbones are shown as cartoon tubes with residues shown as sticks and colored as Extended Data Fig. 7a. Cryo-EM density for the PLLs is shown as a grey transparent surface. Pore facing residues are annotated.

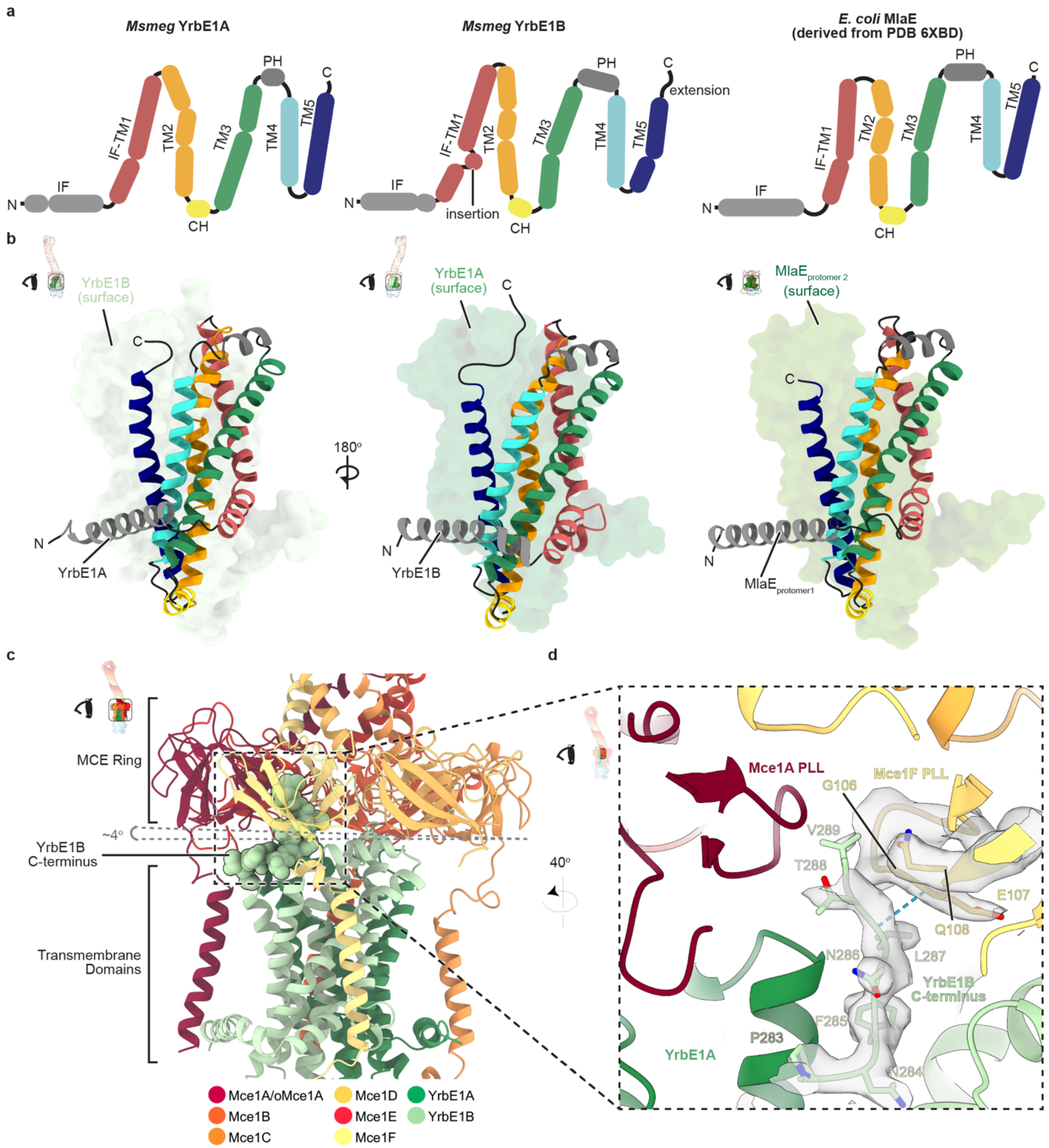


**Extended Data Fig. 8: Structural features of ABC transporter transmembrane domains, YrbE1A and YrbE1B. a**, 2D topology diagram of YrbE1A, Yrbe1B and *E. coli* homolog MlaE (derived from PDB 6XBD[46]). CH, coupling helix; PH, periplasmic helix; IF, interfacial helix; TM, transmembrane helix. **b**, Structures of YrbE1A, YrbE1B and MlaE. YrbE1A and YrbE1B form a heterodimer, while MlaE forms a homodimer. One protomer is shown as a cartoon and the other as a molecular surface in each representation. **c**, View of Mce1 IM complex as indicated by inset on upper-left. Model colored as in the key. The C-terminus of YrbE1B (residues 280-289) is shown as spheres. Grey-dotted lines indicate the MCE ring tilt relative to YrbE1B. **d**, Zoom-in view of region boxed in Extended Data Fig. 8c, oriented as indicated by inset, highlighting interaction between YrbE1B C-terminus and Mce1F PLL. Proteins are shown as cartoon ribbons and colored as Extended Data Fig. 8c. Model is superimposed on cryo-EM density (shown as a transparent grey surface) for the YrbE1B C-terminus (chain J, residues 280-289) and Mce1F PLL (chain F, residues 99-110). Hydrogen bonds are shown as cyan dotted lines.

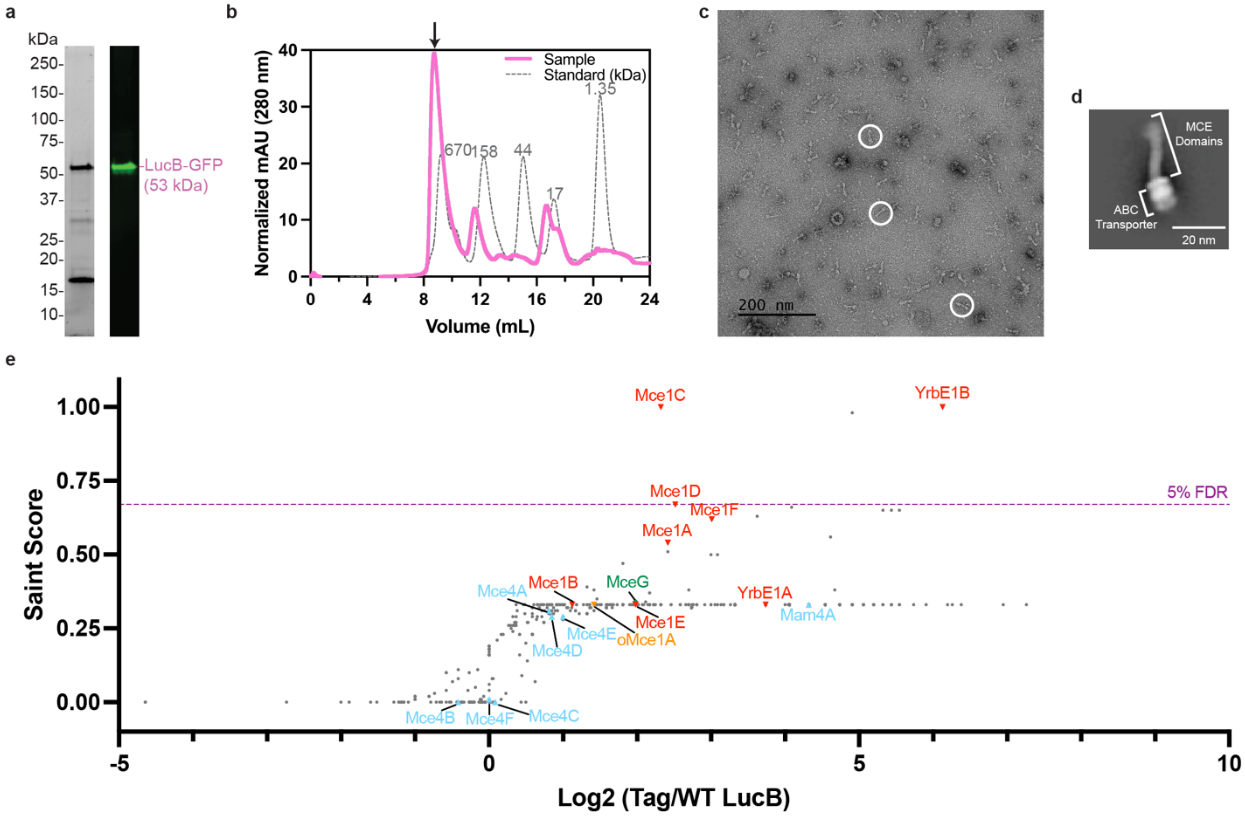


**Extended Data Fig. 9: LucB pulldown, purification and mass spectrometry results. a**, (left) Stain-free SDS-PAGE of LucB-GFP pulldown in n-dodecyl-β-D-maltoside (DDM). (right) Western blot using an anti-GFP antibody against purified LucB-GFP. Band for LucB-GFP indicated by pink label. **b**, Size exclusion chromatogram of LucB-GFP pulldown (pink line) superimposed with protein standards (grey dotted line). Black arrow indicates protein sample further analyzed by negative stain electron microscopy Extended Data Fig. 9c,d and mass spectrometry Extended Data Fig. 9e. **c**, Negative stain electron microscopy micrograph of purified LucB-GFP. Particles of interest, which resemble the Mce1 complex, are circled in white. **d**, 2D class average of 'MCE-like' particles from negative stain electron microscopy of endogenously purified LucB-GFP. Scale bar, 20 nm. **e**, Plot of proteins identified by mass spectrometry that co-purify with LucB-GFP. Each point corresponds to an individual protein plotted by fold change difference after purification of LucB-GFP from *Msmeg* strain harboring tagged LucB versus control wild-type *Msmeg* mc$^2$155 (x-axis) and the probability that a protein is a LucB interactor (SAINT score; y-axis). SAINT score = 1 identifies proteins with the highest probability of being a LucB interactor[59]. SAINT score ≥ 0.67 yielded an FDR (false discovery rate) of ≤ 5% as indicated by the purple dotted line. Proteins related to the mycobacterial MCE systems are highlighted and annotated: Mce1 proteins (red upside-down triangles), Mce4 proteins (sky blue triangles), orphaned MCE proteins (orange upside-down triangle), MceG (green diamond). Proteins outside of these systems are shown as grey dots. Plotted data are from three biological replicates ($n$ = 3).

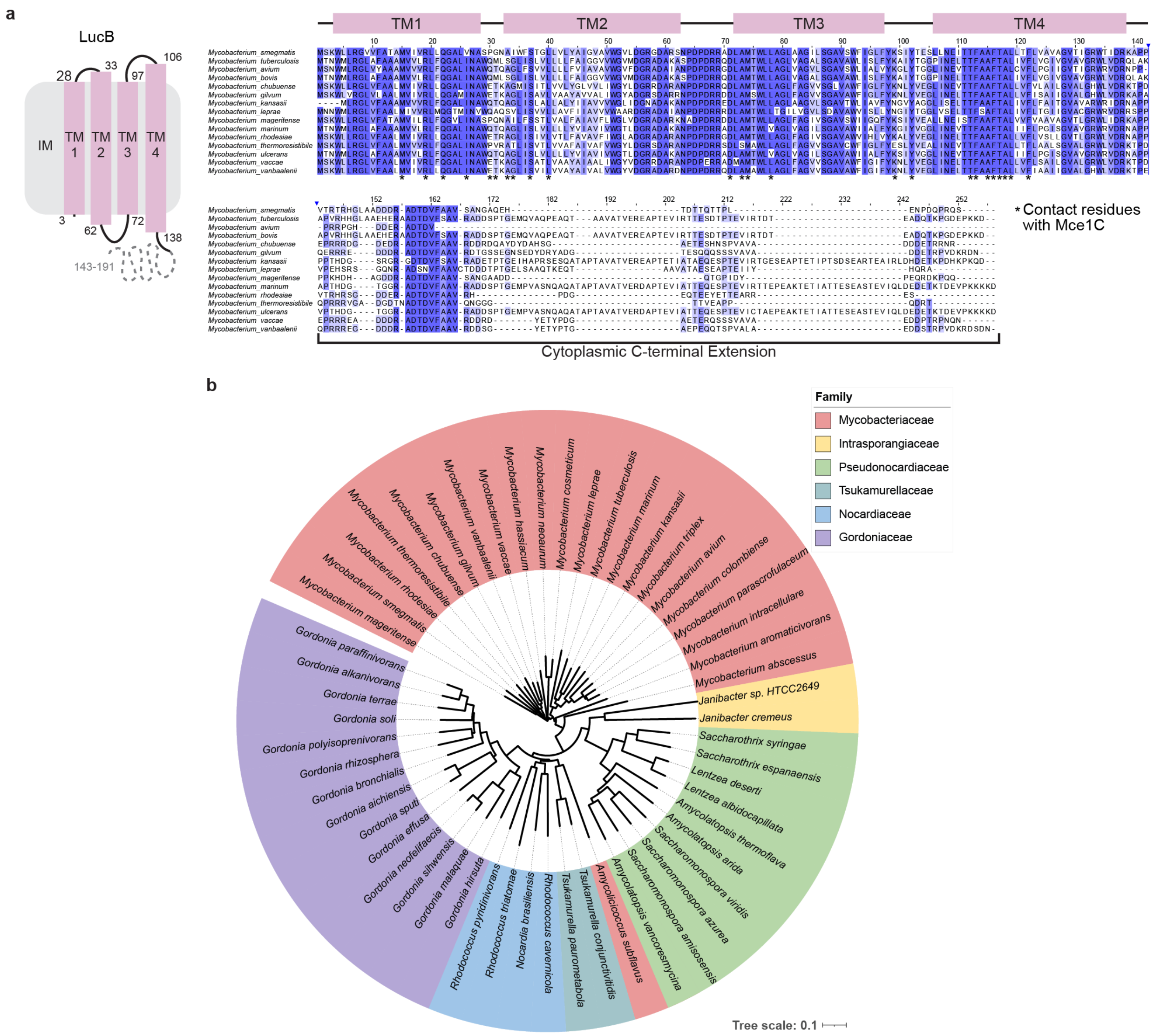


**Extended Data Fig. 10: LucB orthologs are found in other bacteria with MCE systems. a**, (left) 2D topology diagram of LucB. (right) Protein sequence alignment of LucB orthologs from fifteen different mycobacterial species generated using MUSCLE[69] and visualized using JalView[70]. Sequence alignment is colored by percent identity (not identical, white; identical, blue). TMs are depicted above and residues that interact with Mce1C are highlighted below with asterisks. **b**, Circular phylogenetic tree showing spread of orthologs of LucB in other bacteria. LucB orthologs were compiled from eggNOG v5.0[71] and AlphaFold Protein Structure Database[56]. Protein sequences were aligned to generate a phylogenetic tree using MUSCLE[69] and the tree was visualized in ITOL[72]. Leaves indicate individual species and are colored by bacterial family: Mycobacteriaceae (red), Intrasporangiaceae (yellow), Pseudonocardiaceae (green), Tsukamurellaceae (cyan), Nocardiaceae (blue), Gordoniaceae (purple). Tree scale indicated on bottom right.

**Supplementary Table 1 (separate file). Chart of bacterial strains and constructs.**

**Supplementary Table 2 (separate file). Peptide spectral matches for *Msmeg* proteins that co-purified with McеG-GFP and LucB-GFP.**

**Supplementary Table 3 (separate file). SAINT scores for *Msmeg* proteins that co-purified with MceG-GFP.**

**Supplementary Table 4 (separate file). Cryo-EM data collection, refinement, and validation statistics.**

**Supplementary Table 5 (separate file). SAINT scores for *Msmeg* proteins that co-purified with LucB-GFP.**

**Supplementary Table 6 (separate file). AlphaFold2 predictions used for cryo-EM modeling.**

**Supplementary Information Figure 1 (separate file). Uncropped gel and blot from Figure 1e.**

**Supplementary Information Figure 2 (separate file). Uncropped gel and blot from Extended Data Figure 9a.**

**Supplementary Video 1 (separate file). Overview of Mce1 structure.**

**Supplementary Video 2 (separate file). Model for Mce1-mediated transport.**
