## Supplementary Table 1 for "Structure of an endogenous mycobacterial MCE lipid transporter"

**Supplementary Table 1. Chart of bacterial strains and constructs.**

| Reagent or Resource | Description | Source | Identifier |
| --- | --- | --- | --- |
| Experimental Models: Organisms/Strains | | | |
| *E. coli* TOP10 |  | Invitrogen | C404010 |
| mc^2^155 | Highly transformable variant of *M. smegmatis* | Snapper et al., 1990 |  |
| bBEL591 | mc^2^155 *mceG-*3C-eGFP-4xGly-Tev-Flag-His6 | This study |  |
| bBEL594 | mc^2^155 Δ*mceG* | This study |  |
| bBEL595 | mc^2^155 *lucB*-3C-eGFP-4xGly-Tev-Flag-His6 | This study |  |
| Recombinant DNA | | | |
| pKM444 | For performing ORBIT - expresses Che9c phage RecT and Bxb1 phage Integrase | Murphy et al., 2018 | Addgene #108319 |
| pKM461 | For performing ORBIT - expresses Che9c phage RecT and Bxb1 phage Integrase, and and contains SacB | Murphy et al., 2018 | Addgene #108320 |
| pKM464 | ORBIT Integrating plasmid to delete target gene. | Murphy et al., 2018 | Addgene #108322 |
| pKM468-EGFP | ORBIT integrating plasmid for C-terminal tagging with EGFP. | Murphy et al., 2018 | Addgene #108434 |
| pBEL2108 | pKM468-EGFP derivative containing a 3C protease cleavage site upstream of the eGFP tag | This study |  |
| pBEL1782 | pMV261 with a Zeocin resistance cassette (Zeo^R^) | Gift from Jeffery Cox (UC Berkeley) |  |
| pBEL2759 | pMV261Zeo^R^-*mceG* *(MSMEG_1366)* | This study |  |
| pTP396 | Expresses His14-Biotin_acceptor_peptide-SUMOEu1-anti-GFP_nanobody | Pleiner et al., 2020 | Addgene# 149336 |
| pTP264 | Expresses Biotin ligase BirA | Pleiner et al., 2020 | Addgene# 149334 |
| pAV286 | Expresses SENP_EuB protease | Pleiner et al., 2020 | Addgene# 149333 |
| pBEL2719 | pMV261Zeo^R^-*mceG (MSMEG_1366)* Y178A | This study |  |
| pBEL2713 | pMV261Zeo^R^-*mceG (MSMEG_1366)* 𝚫242-360 | This study |  |
