## Supplementary Table 4 for "Structure of an endogenous mycobacterial MCE lipid transporter"

**Supplementary Table 4. Cryo-EM data collection, refinement, and validation statistics.**

|  | Map0a | Map0b | Map0c | Map0d | Map0e | Map0 composite |
| --- | --- | --- | --- | --- | --- | --- |
|  | (EMD-  29228) | (EMD-  29229) | (EMD-  29230) | (EMD-  29231) | (EMD-  29232) | (EMD-29025) |
|  |  |  |  |  |  | (PDB ID 8FEF) |
| **Data collection and processing** | | | | | | |
| Dataset 1 |  |  |  |  |  |  |
| Microscope | Krios G3 (PNCC Krios #2) | Krios G3 (PNCC Krios #2) | Krios G3 (PNCC Krios #2) | Krios G3 (PNCC Krios #2) | Krios G3 (PNCC Krios #2) | Krios G3 (PNCC Krios #2) |
| Voltage (keV) | 300 | 300 | 300 | 300 | 300 | 300 |
| Camera | K3 Bio Continuum | K3 Bio Continuum | K3 Bio Continuum | K3 Bio Continuum | K3 Bio Continuum | K3 Bio Continuum |
| Magnification | 105,000x | 105,000x | 105,000x | 105,000x | 105,000x | 105,000x |
| Nominal Pixel size (Å/pixel) | 0.8255 | 0.8255 | 0.8255 | 0.8255 | 0.8255 | 0.8255 |
| Total electron exposure  (e–/Å2) | 60 | 60 | 60 | 60 | 60 | 60 |
| Number of frames (no.) | 60 | 60 | 60 | 60 | 60 | 60 |
| Defocus range (μm) | -0.8 to  -2.4 | -0.8 to  -2.4 | -0.8 to  -2.4 | -0.8 to  -2.4 | -0.8 to  -2.4 | -0.8 to  -2.4 |
| Automation software | Serial-EM | Serial-EM | Serial-EM | Serial-EM | Serial-EM | Serial-EM |
| Energy filter slit width (eV) | 20 | 20 | 20 | 20 | 20 | 20 |
| Micrographs collected (no.) | 21,915 | 21,915 | 21,915 | 21,915 | 21,915 | 21,915 |
| Total extracted particles (no.) | 1,820,584 | 1,820,584 | 1,820,584 | 1,820,584 | 1,820,584 | 1,820,584 |
| Dataset 2 |  |  |  |  |  |  |
| Microscope | Krios G3 (PNCC Krios #2) | Krios G3 (PNCC Krios #2) | Krios G3 (PNCC Krios #2) | Krios G3 (PNCC Krios #2) | Krios G3 (PNCC Krios #2) | Krios G3 (PNCC Krios #2) |
| Voltage (keV) | 300 | 300 | 300 | 300 | 300 | 300 |
| Camera | K3 Bio Continuum | K3 Bio Continuum | K3 Bio Continuum | K3 Bio Continuum | K3 Bio Continuum | K3 Bio Continuum |
| Magnification | 105,000 | 105,000 | 105,000 | 105,000 | 105,000 | 105,000 |
| Pixel size at detector (Å/pixel) | 0.8255 | 0.8255 | 0.8255 | 0.8255 | 0.8255 | 0.8255 |
| Total electron exposure  (e–/Å2) | 60 | 60 | 60 | 60 | 60 | 60 |
| Number of frames (no.) | 60 | 60 | 60 | 60 | 60 | 60 |
| Defocus range (μm) | -0.8 to  -2.4 | -0.8 to  -2.4 | -0.8 to  -2.4 | -0.8 to  -2.4 | -0.8 to  -2.4 | -0.8 to  -2.4 |
| Automation software | Serial-EM | Serial-EM | Serial-EM | Serial-EM | Serial-EM | Serial-EM |
| Tilt angle (^o^) | -30 | -30 | -30 | -30 | -30 | -30 |
| Energy filter slit width (eV) | 20 | 20 | 20 | 20 | 20 | 20 |
| Micrographs collected (no.) | 22,010 | 22,010 | 22,010 | 22,010 | 22,010 | 22,010 |
| Total extracted particles (no.) | 1,048,639 | 1,048,639 | 1,048,639 | 1,048,639 | 1,048,639 | 1,048,639 |
| Dataset 1 and 2 (combined) | | | | | | |
| Box size (px) | 360 | 360 | 360 | 360 | 640 | 640 |
| Final particle images (no.) | 340,238 | 341,490 | 334,050 | 330,104 | 305,172 | 305,172 |
| Symmetry | C1 | C1 | C1 | C1 | C1 | C1 |
| Map resolution (Å) |  |  |  |  |  |  |
| FSC threshold 0.5 | 3.4 | 3.1 | 3.2 | 3.1 | 3.2 | 3.2 |
| FSC threshold 0.143 | 2.91 | 2.73 | 2.75 | 3.00 | 2.92 | 2.71 |
| Map resolution range (Å) | 2.6-6.3 | 2.4-5.6 | 2.5-6.0 | 2.7-7.3 | 2.8-7.2 | 2.3-4.9 |
| Map sharpening B factor (Å2) | 98.3 | 88.7 | 91.1 | 102.0 | 76.6 |  |
| Sphericity | 0.821 | 0.869 | 0.853 | 0.853 |  |  |
| **Model composition, refinement, validation** | | | | | | |
| Initial models used (PDB code) |  |  |  |  |  | AlphaFold |
| Model composition |  |  |  |  |  |  |
| Chains |  |  |  |  |  | 11 |
| Non-hydrogen atoms |  |  |  |  |  | 25,533 |
| Protein residues |  |  |  |  |  | 3,292 |
| Ligands |  |  |  |  |  | UNL: 31 |
| Refinement |  |  |  |  |  |  |
| CC (mask) |  |  |  |  |  | 0.80 |
| CC (box) |  |  |  |  |  | 0.62 |
| CC (peaks) |  |  |  |  |  | 0.58 |
| CC (volume) |  |  |  |  |  | 0.79 |
| Mean CC for ligands |  |  |  |  |  | 0.70 |
| *B* factors (Å^2^) |  |  |  |  |  |  |
| Protein |  |  |  |  |  | 74.65 |
| Ligand |  |  |  |  |  | 70.79 |
| R.M.S. deviations |  |  |  |  |  |  |
| Bond lengths (Å) |  |  |  |  |  | 0.005 |
| Bond angles (°) |  |  |  |  |  | 1.025 |
| Validation |  |  |  |  |  |  |
| MolProbity score |  |  |  |  |  | 1.48 |
| CaBLAM outliers (%) |  |  |  |  |  | 1.36 |
| Clashscore |  |  |  |  |  | 6.44 |
| Poor rotamers (%) |  |  |  |  |  | 0.85 |
| Cβ outliers (%) |  |  |  |  |  | 0.00 |
| EMRinger score |  |  |  |  |  | 3.46 |
| Ramachandran plot |  |  |  |  |  |  |
| Favored (%) |  |  |  |  |  | 97.37 |
| Allowed (%) |  |  |  |  |  | 2.54 |
| Disallowed (%) |  |  |  |  |  | 0.09 |
| Rama-Z |  |  |  |  |  |  |
| whole |  |  |  |  |  | 1.24 |
| helix |  |  |  |  |  | 1.86 |
| sheet |  |  |  |  |  | 0.37 |
| loop |  |  |  |  |  | -0.58 |

|  | Map1a | Map1b | Map1c | Map1d | Map1e | Map1 composite |
| --- | --- | --- | --- | --- | --- | --- |
|  | (EMD-  29233) | (EMD-  29234) | (EMD-  29235) | (EMD-  29236) | (EMD-  29237) | (EMD-29023) |
|  |  |  |  |  |  | (PDB ID 8FED) |
| **Data collection and processing** | | | | | | |
| Dataset 1 |  |  |  |  |  |  |
| Microscope | Krios G3 (PNCC Krios #2) | Krios G3 (PNCC Krios #2) | Krios G3 (PNCC Krios #2) | Krios G3 (PNCC Krios #2) | Krios G3 (PNCC Krios #2) | Krios G3 (PNCC Krios #2) |
| Voltage (keV) | 300 | 300 | 300 | 300 | 300 | 300 |
| Camera | K3 Bio Continuum | K3 Bio Continuum | K3 Bio Continuum | K3 Bio Continuum | K3 Bio Continuum | K3 Bio Continuum |
| Magnification | 105,000x | 105,000x | 105,000x | 105,000x | 105,000x | 105,000x |
| Nominal Pixel size (Å/pixel) | 0.8255 | 0.8255 | 0.8255 | 0.8255 | 0.8255 | 0.8255 |
| Total electron exposure (e–/Å2) | 60 | 60 | 60 | 60 | 60 | 60 |
| Number of frames (no.) | 60 | 60 | 60 | 60 | 60 | 60 |
| Defocus range (μm) | -0.8 to  -2.4 | -0.8 to  -2.4 | -0.8 to  -2.4 | -0.8 to  -2.4 | -0.8 to  -2.4 | -0.8 to  -2.4 |
| Automation software | Serial-EM | Serial-EM | Serial-EM | Serial-EM | Serial-EM | Serial-EM |
| Energy filter slit width (eV) | 20 | 20 | 20 | 20 | 20 | 20 |
| Micrographs collected (no.) | 21,915 | 21,915 | 21,915 | 21,915 | 21,915 | 21,915 |
| Total extracted particles (no.) | 1,820,584 | 1,820,584 | 1,820,584 | 1,820,584 | 1,820,584 | 1,820,584 |
| Dataset 2 |  |  |  |  |  |  |
| Microscope | Krios G3 (PNCC Krios #2) | Krios G3 (PNCC Krios #2) | Krios G3 (PNCC Krios #2) | Krios G3 (PNCC Krios #2) | Krios G3 (PNCC Krios #2) | Krios G3 (PNCC Krios #2) |
| Voltage (keV) | 300 | 300 | 300 | 300 | 300 | 300 |
| Camera | K3 Bio Continuum | K3 Bio Continuum | K3 Bio Continuum | K3 Bio Continuum | K3 Bio Continuum | K3 Bio Continuum |
| Magnification | 105,000 | 105,000 | 105,000 | 105,000 | 105,000 | 105,000 |
| Pixel size at detector (Å/pixel) | 0.8255 | 0.8255 | 0.8255 | 0.8255 | 0.8255 | 0.8255 |
| Total electron exposure  (e–/Å2) | 60 | 60 | 60 | 60 | 60 | 60 |
| Number of frames (no.) | 60 | 60 | 60 | 60 | 60 | 60 |
| Defocus range (μm) | -0.8 to  -2.4 | -0.8 to  -2.4 | -0.8 to  -2.4 | -0.8 to  -2.4 | -0.8 to  -2.4 | -0.8 to  -2.4 |
| Automation software | Serial-EM | Serial-EM | Serial-EM | Serial-EM | Serial-EM | Serial-EM |
| Tilt angle (^o^) | -30 | -30 | -30 | -30 | -30 | -30 |
| Energy filter slit width (eV) | 20 | 20 | 20 | 20 | 20 | 20 |
| Micrographs collected (no.) | 22,010 | 22,010 | 22,010 | 22,010 | 22,010 | 22,010 |
| Total extracted particles (no.) | 1,048,639 | 1,048,639 | 1,048,639 | 1,048,639 | 1,048,639 | 1,048,639 |
| **Dataset 1 and 2 (combined)** | | | | | | |
| Box size (px) | 360 | 360 | 360 | 360 | 640 | 640 |
| Final particle images (no.) | 161,434 | 162,004 | 158,508 | 156,741 | 144,822 | 144,822 |
| Symmetry | C1 | C1 | C1 | C1 | C1 | C1 |
| Map resolution (Å) |  |  |  |  |  |  |
| FSC threshold 0.5 | 3.6 | 3.3 | 3.3 | 3.8 | 3.5 | 3.1 |
| FSC threshold 0.143 | 3.05 | 2.89 | 2.97 | 3.16 | 3.04 | 2.76 |
| Map resolution range (Å) | 2.7-6.7 | 2.6-5.9 | 2.7-6.4 | 2.8-7.7 | 2.8-9.6 | 2.3-5.5 |
| Map sharpening B factor (Å2) | 89.2 | 79.4 | 80.7 | 93.9 | 63.3 |  |
| Sphericity | 0.677 | 0.771 | 0.727 | 0.71 |  |  |
| **Model composition, refinement, validation** | | | | | | |
| Initial models used (PDB code) |  |  |  |  |  | AlphaFold |
| Model composition |  |  |  |  |  |  |
| Chains |  |  |  |  |  | 12 |
| Non-hydrogen atoms |  |  |  |  |  | 26,565 |
| Protein residues |  |  |  |  |  | 3,427 |
| Ligands |  |  |  |  |  | UNL: 31 |
| Refinement |  |  |  |  |  |  |
| CC (mask) |  |  |  |  |  | 0.77 |
| CC (box) |  |  |  |  |  | 0.63 |
| CC (peaks) |  |  |  |  |  | 0.56 |
| CC (volume) |  |  |  |  |  | 0.76 |
| Mean CC for ligands |  |  |  |  |  | 0.66 |
| *B* factors (Å^2^) |  |  |  |  |  |  |
| Protein |  |  |  |  |  | 82.16 |
| Ligand |  |  |  |  |  | 75.06 |
| R.M.S. deviations |  |  |  |  |  |  |
| Bond lengths (Å) |  |  |  |  |  | 0.004 |
| Bond angles (°) |  |  |  |  |  | 0.985 |
| Validation |  |  |  |  |  |  |
| MolProbity score |  |  |  |  |  | 1.46 |
| CaBLAM outliers (%) |  |  |  |  |  | 1.49 |
| Clashscore |  |  |  |  |  | 5.69 |
| Poor rotamers (%) |  |  |  |  |  | 0.78 |
| Cβ outliers (%) |  |  |  |  |  | 0.00 |
| EMRinger score |  |  |  |  |  | 2.84 |
| Ramachandran plot |  |  |  |  |  |  |
| Favored (%) |  |  |  |  |  | 97.14 |
| Allowed (%) |  |  |  |  |  | 2.83 |
| Disallowed (%) |  |  |  |  |  | 0.03 |
| Rama-Z |  |  |  |  |  |  |
| whole |  |  |  |  |  | 1.44 |
| helix |  |  |  |  |  | 2.01 |
| sheet |  |  |  |  |  | 0.15 |
| loop |  |  |  |  |  | -0.49 |

|  | Map2a | Map2b | Map2c | Map2d | Map2e | Map2 composite |
| --- | --- | --- | --- | --- | --- | --- |
|  | (EMD-  29238) | (EMD-  29239) | (EMD-  29240) | (EMD-  29241) | (EMD-  29242) | (EMD-29024) |
|  |  |  |  |  |  | (PDB ID 8FEE) |
| **Data collection and processing** | | | | | | |
| Dataset 1 |  |  |  |  |  |  |
| Microscope | Krios G3 (PNCC Krios #2) | Krios G3 (PNCC Krios #2) | Krios G3 (PNCC Krios #2) | Krios G3 (PNCC Krios #2) | Krios G3 (PNCC Krios #2) | Krios G3 (PNCC Krios #2) |
| Voltage (keV) | 300 | 300 | 300 | 300 | 300 | 300 |
| Camera | K3 Bio Continuum | K3 Bio Continuum | K3 Bio Continuum | K3 Bio Continuum | K3 Bio Continuum | K3 Bio Continuum |
| Magnification | 105,000x | 105,000x | 105,000x | 105,000x | 105,000x | 105,000x |
| Nominal Pixel size (Å/pixel) | 0.8255 | 0.8255 | 0.8255 | 0.8255 | 0.8255 | 0.8255 |
| Total electron exposure  (e–/Å2) | 60 | 60 | 60 | 60 | 60 | 60 |
| Number of frames (no.) | 60 | 60 | 60 | 60 | 60 | 60 |
| Defocus range (μm) | -0.8 to  -2.4 | -0.8 to  -2.4 | -0.8 to  -2.4 | -0.8 to  -2.4 | -0.8 to  -2.4 | -0.8 to  -2.4 |
| Automation software | Serial-EM | Serial-EM | Serial-EM | Serial-EM | Serial-EM | Serial-EM |
| Energy filter slit width (eV) | 20 | 20 | 20 | 20 | 20 | 20 |
| Micrographs collected (no.) | 21,915 | 21,915 | 21,915 | 21,915 | 21,915 | 21,915 |
| Total extracted particles (no.) | 1,820,584 | 1,820,584 | 1,820,584 | 1,820,584 | 1,820,584 | 1,820,584 |
| Dataset 2 |  |  |  |  |  |  |
| Microscope | Krios G3 (PNCC Krios #2) | Krios G3 (PNCC Krios #2) | Krios G3 (PNCC Krios #2) | Krios G3 (PNCC Krios #2) | Krios G3 (PNCC Krios #2) | Krios G3 (PNCC Krios #2) |
| Voltage (keV) | 300 | 300 | 300 | 300 | 300 | 300 |
| Camera | K3 Bio Continuum | K3 Bio Continuum | K3 Bio Continuum | K3 Bio Continuum | K3 Bio Continuum | K3 Bio Continuum |
| Magnification | 105,000 | 105,000 | 105,000 | 105,000 | 105,000 | 105,000 |
| Pixel size at detector (Å/pixel) | 0.8255 | 0.8255 | 0.8255 | 0.8255 | 0.8255 | 0.8255 |
| Total electron exposure  (e–/Å2) | 60 | 60 | 60 | 60 | 60 | 60 |
| Number of frames (no.) | 60 | 60 | 60 | 60 | 60 | 60 |
| Defocus range (μm) | -0.8 to  -2.4 | -0.8 to  -2.4 | -0.8 to  -2.4 | -0.8 to  -2.4 | -0.8 to  -2.4 | -0.8 to  -2.4 |
| Automation software | Serial-EM | Serial-EM | Serial-EM | Serial-EM | Serial-EM | Serial-EM |
| Tilt angle (^o^) | -30 | -30 | -30 | -30 | -30 | -30 |
| Energy filter slit width (eV) | 20 | 20 | 20 | 20 | 20 | 20 |
| Micrographs collected (no.) | 22,010 | 22,010 | 22,010 | 22,010 | 22,010 | 22,010 |
| Total extracted particles (no.) | 1,048,639 | 1,048,639 | 1,048,639 | 1,048,639 | 1,048,639 | 1,048,639 |
| Dataset 1 and 2 (combined) | | | | | | |
| Box size (px) | 360 | 360 | 360 | 360 | 640 | 640 |
| Final particle images (no.) | 178,844 | 179,480 | 175,490 | 173,315 | 160,443 | 160,443 |
| Symmetry | C1 | C1 | C1 | C1 | C1 | C1 |
| Map resolution (Å) |  |  |  |  |  |  |
| FSC threshold 0.5 | 3.7 | 3.5 | 3.6 | 3.8 | 3.6 | 3.3 |
| FSC threshold 0.143 | 3.13 | 2.99 | 3.06 | 3.19 | 3.19 | 2.90 |
| Map resolution range (Å) | 2.8-6.4 | 2.6-6.4 | 2.7-6.5 | 2.8-7.6 | 2.8-8.0 | 3.0-5.7 |
| Map sharpening B factor (Å2) | 95.4 | 81.1 | 82 | 91.6 | 67.3 |  |
| Sphericity | 0.766 | 0.772 | 0.745 | 0.729 |  |  |
| **Model composition, refinement, validation** | | | | | | |
| Initial models used (PDB code) |  |  |  |  |  | AlphaFold |
| Model composition |  |  |  |  |  |  |
| Chains |  |  |  |  |  | 11 |
| Non-hydrogen atoms |  |  |  |  |  | 25,218 |
| Protein residues |  |  |  |  |  | 3,248 |
| Ligands |  |  |  |  |  | UNL: 31 |
| Refinement |  |  |  |  |  |  |
| CC (mask) |  |  |  |  |  | 0.76 |
| CC (box) |  |  |  |  |  | 0.65 |
| CC (peaks) |  |  |  |  |  | 0.58 |
| CC (volume) |  |  |  |  |  | 0.74 |
| Mean CC for ligands |  |  |  |  |  | 0.66 |
| *B* factors (Å^2^) |  |  |  |  |  |  |
| Protein |  |  |  |  |  | 77.69 |
| Ligand |  |  |  |  |  | 64.88 |
| R.M.S. deviations |  |  |  |  |  |  |
| Bond lengths (Å) |  |  |  |  |  | 0.004 |
| Bond angles (°) |  |  |  |  |  | 1.015 |
| Validation |  |  |  |  |  |  |
| MolProbity score |  |  |  |  |  | 1.62 |
| CaBLAM outliers (%) |  |  |  |  |  | 1.47 |
| Clashscore |  |  |  |  |  | 5.97 |
| Poor rotamers (%) |  |  |  |  |  | 1.12 |
| Cβ outliers (%) |  |  |  |  |  | 0.00 |
| EMRinger score |  |  |  |  |  | 2.45 |
| Ramachandran plot |  |  |  |  |  |  |
| Favored (%) |  |  |  |  |  | 96.24 |
| Allowed (%) |  |  |  |  |  | 3.73 |
| Disallowed (%) |  |  |  |  |  | 0.03 |
| Rama-Z |  |  |  |  |  |  |
| whole |  |  |  |  |  | 0.97 |
| helix |  |  |  |  |  | 1.64 |
| sheet |  |  |  |  |  | -0.09 |
| loop |  |  |  |  |  | -0.58 |
