## Supplementary Table 6 for "Structure of an endogenous mycobacterial MCE lipid transporter"

**Supplementary Table 6. AlphaFold2 predictions used for cryo-EM modeling.**

| Model | AlphaFold Prediction | Server | Usage |
| --- | --- | --- | --- |
| AFpdb1 | MceG [MSMEG_1366] | ColabFold | Access subunit identity and placement in cryo-EM map |
| AFpdb2 | YrbE1A [MSMEG_0132] | ColabFold | Access subunit identity and placement in cryo-EM map |
| AFpdb3 | YrbE1B [MSMEG_0133] | ColabFold | Access subunit identity and placement in cryo-EM map |
| AFpdb4 | Mce1A [MSMEG_0134] | ColabFold | Access subunit identity and placement in cryo-EM map |
| AFpdb5 | Mce1B [MSMEG_0135] | ColabFold | Access subunit identity and placement in cryo-EM map |
| AFpdb6 | Mce1C [MSMEG_0136] | ColabFold | Access subunit identity and placement in cryo-EM map |
| AFpdb7 | Mce1D [MSMEG_0137] | ColabFold | Access subunit identity and placement in cryo-EM map |
| AFpdb8 | Mce1E [MSMEG_138] | ColabFold | Access subunit identity and placement in cryo-EM map |
| AFpdb9 | Mce1F [MSMEG_139] | ColabFold | Access subunit identity and placement in cryo-EM map |
| AFpdb10 | YrbE4A [MSMEG_5902] | ColabFold | Access subunit identity and placement in cryo-EM map |
| AFpdb11 | YrbE4B [MSMEG_5901] | ColabFold | Access subunit identity and placement in cryo-EM map |
| AFpdb12 | Mce4A [MSMEG_5900] | ColabFold | Access subunit identity and placement in cryo-EM map |
| AFpdb13 | Mce4B [MSMEG_5899] | ColabFold | Access subunit identity and placement in cryo-EM map |
| AFpdb14 | Mce4C [MSMEG_5898] | ColabFold | Access subunit identity and placement in cryo-EM map |
| AFpdb15 | Mce4D [MSMEG_5897] | ColabFold | Access subunit identity and placement in cryo-EM map |
| AFpdb16 | Mce4E [MSMEG_5896] | ColabFold | Access subunit identity and placement in cryo-EM map |
| AFpdb17 | Mce4F [MSMEG_5895] | ColabFold | Access subunit identity and placement in cryo-EM map |
| AFpdb18 | oMce1A [MSMEG_6540] | ColabFold | Access subunit identity and placement in cryo-EM map |
| AFpdb19 | MceG (1-360)/MceG (1-360)/YrbE1A/YrbE1B) | ColabFold | Initial model building into cryo-EM map |
| AFpdb20 | YrbE1A/YrbE1B/Mce1A (1-409)/Mce1B  (1-343)/Mce1C (1-524)/Mce1D (1-547)/Mce1E (1-390)/Mce1F (1-518)) | COSMIC2 | Initial model building into cryo-EM map |
| AFpdb21 | (Mce1A (151-409)/Mce1B (135-343)/Mce1C (135-524)/Mce1D (146-547)/Mce1E (152-390)/Mce1F (136-518)) | COSMIC2 | Initial model building into cryo-EM map |
| AFpdb22 | LucB [MSMEG_3032] | ColabFold | Initial model building into cryo-EM map |
