## Supplementary Information Figure 1 for "Structure of an endogenous mycobacterial MCE lipid transporter"

**Fig. 1e**

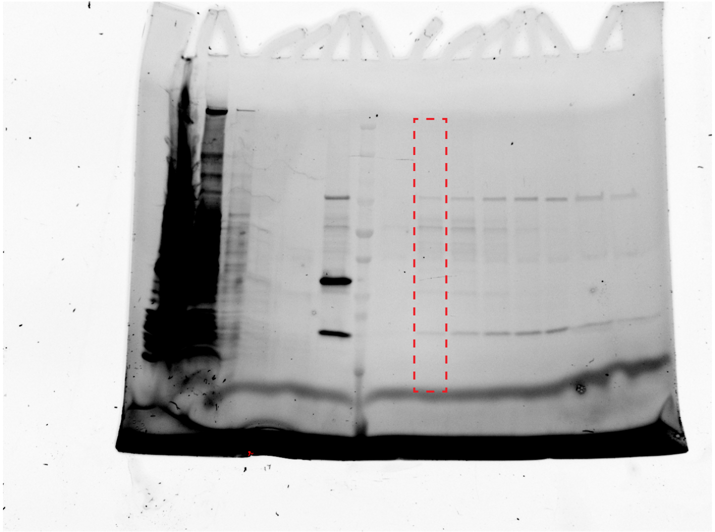

MceG-GFP Purification (Stain-free SDS-PAGE gel)

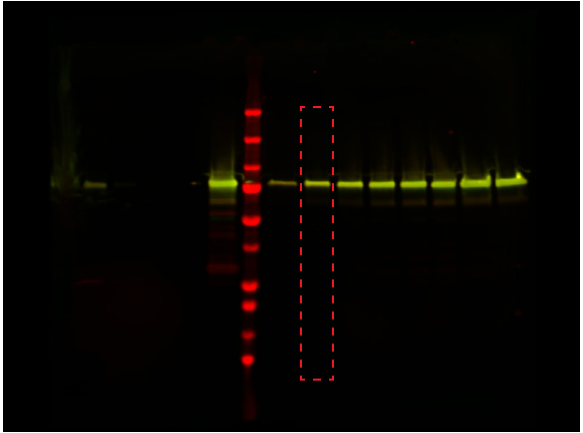

MceG-GFP Purification (Western-Blot, 700 & 800 channels merged)
