## Supplementary Information Figure 2 for "Structure of an endogenous mycobacterial MCE lipid transporter"

**Extended Data Fig. 9a**

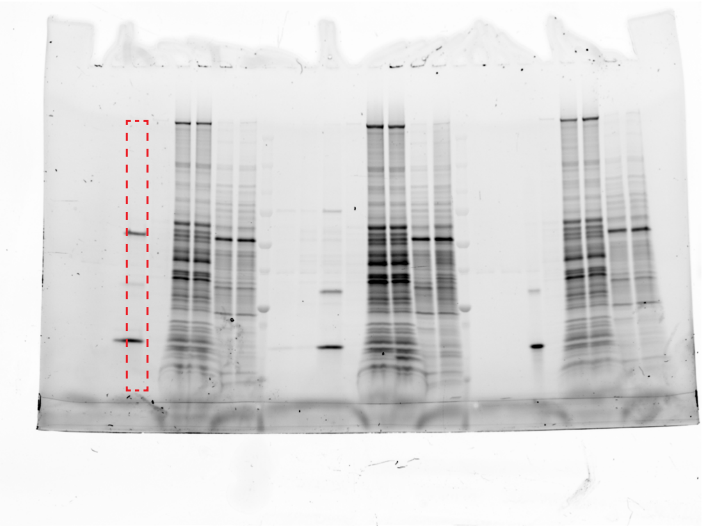

LucB-GFP Purification (Stain-free SDS-PAGE gel)

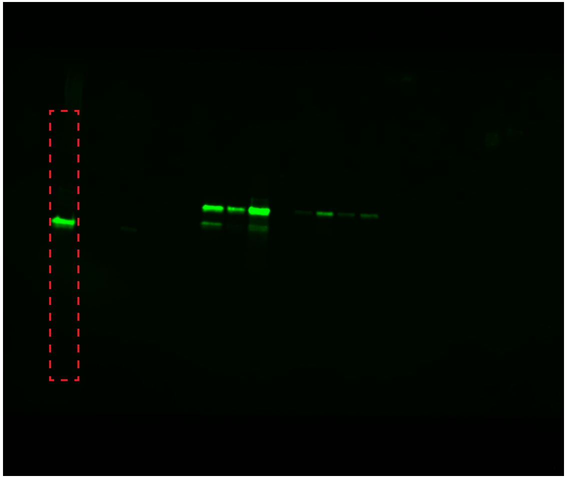

LucB-GFP Purification (Western-Blot, 800 channel)

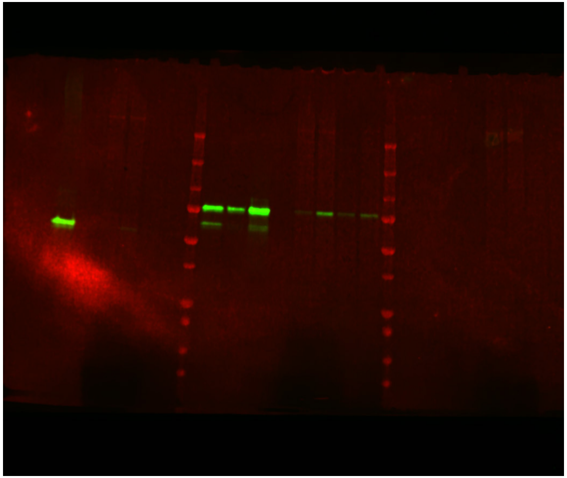

MceG-GFP Purification (Western-Blot, 700 & 800 channel merged)
